# *INTS13* Mutations Causing a Developmental Ciliopathy Disrupt Integrator Complex Assembly

**DOI:** 10.1101/2020.07.20.209130

**Authors:** Lauren G. Mascibroda, Mohammad Shboul, Nathan D. Elrod, Laurence Colleaux, Hanan Hamamy, Kai-Lieh Huang, Natoya Peart, Moirangthem Kiran Singh, Hane Lee, Barry Merriman, Jeanne N. Jodoin, Laura A. Lee, Raja Fathalla, Baeth Al-Rawashdeh, Osama Ababneh, Mohammad El-Khateeb, Nathalie Escande-Beillard, Stanley F. Nelson, Yixuan Wu, Liang Tong, Linda J. Kenney, William K. Russell, Jeanne Amiel, Bruno Reversade, Eric J. Wagner

**Affiliations:** Department of Biochemistry and Molecular Biology, The University of Texas Medical Branch at Galveston, Galveston TX 77550, USA; Department of Medical Laboratory Sciences, Jordan University of Science and Technology, Irbid, Jordan; Inserm UMR 1163, Institut Imagine, 24 Boulevard du Montparnasse, 75015 Paris, France; Department of Genetic Medicine and Development, University Hospital, Geneva, Switzerland; Department of Pathology and Laboratory Medicine, Department of Human Genetics, David Geffen School of Medicine, University of California Los Angeles, Los Angeles CA 90095, USA; Department of Cell and Developmental Biology, Vanderbilt University Medical Center, Nashville, TN 37232, USA; Department of Pathology, Microbiology and Immunology, Vanderbilt University Medical Center, Nashville, TN 37232, USA; Faculty of Medicine, Hospital of the University of Jordan, University of Jordan, Jordan; National Center for Diabetes, Endocrinology and Genetics, Jordan; Department of Biological Sciences, Columbia University, New York, NY 10027, USA; Service de Génétique, Institut Imagine, 24 Boulevard du Montparnasse 75015, Paris, France; Department of Paediatrics, School of Medicine, NUS, Singapore; Department of Medical Genetics, KOÇ University, Istanbul, Turkey; Genome Institute of Singapore, A*STAR, 137673, Singapore; Institute of Molecular and Cell Biology, A*STAR, Singapore

## Abstract

Oral-facial-digital syndromes (OFD) are a heterogeneous group of congenital disorders characterized by malformations of the face and oral cavity, and digit anomalies. To date, mutations in 12 ciliary-related genes have been identified that cause several types of OFD, suggesting that OFDs constitute a subgroup of developmental ciliopathies. Through homozygosity mapping and exome sequencing of two families with variable OFD type 2, we identified distinct germline mutations in *INTS13*, a subunit of the Integrator complex. This 14-component complex associates with RNAPII and can cleave nascent RNA to modulate gene expression. We determined that INTS13 utilizes a discrete domain within its C-terminus to bind the Integrator cleavage module, which is disrupted by the identified germline *INTS13* mutations. Depletion of *INTS13* disrupts ciliogenesis in human cultured cells and causes dysregulation of a broad collection of ciliary genes. Accordingly, its knockdown in *Xenopus* embryos lead to motile cilia anomalies. Altogether, we show that mutations in *INTS13* cause an autosomal recessive ciliopathy, which reveals key interactions within Integrator components.

## INTRODUCTION

Ciliopathies encompass a large and expanding group of human genetic disorders caused by mutations in genes either encoding ciliary proteins or that participate indirectly in the formation and function of cilia^1, 2, 3, 4^. There have been several hundred diseases identified as ciliopathies that range from renal cystic disease^5^ and isolated blindness^6^ to multiple organ disorders such as a Bardet-Biedl syndrome (BBS)^7^ or Meckel Gruber syndrome (MKS)^8^. The clinical manifestations associated with ciliopathies are broad and highly variable and include: retinal degeneration, specific nervous system malformations, hepatic, kidney and pancreatic cysts, heterotaxia, polydactyly, encephalocele, hydrocephalus, hearing loss, anosmia, intellectual disability, developmental delay, skeletal malformation, craniofacial anomalies, respiratory function impairment, infertility and obesity^4, 9, 10, 11, 12, 13^.

Among ciliopathies, oral-facial-digital syndromes (OFDs) are a group of clinically and genetically heterogeneous developmental disorders characterized by defects in the development of the face and oral cavity along with digit anomalies. Symptoms associated with OFDs can also be found in other organs, which has led to the classification of at least 14 different types of OFD^14, 15^. To date, 12 genes have been identified to be mutated in OFD syndromes and the encoded proteins have been implicated in primary cilia biogenesis and function. The phenotypic spectrum of OFD syndromes can overlap with other known ciliopathies, and mutations in OFD genes can also cause other types of ciliopathies^16, 17, 18, 19, 20, 21, 22^. Of the 12 known genes involved in OFDs, the majority encode structural components of the primary cilium, such as *OFD1*^23^ and *TMEM231*^24^. However, not all OFD-causing gene mutations such as those in the RNA helicase DDX59^19^ localize to the primary cilium, and are thus thought to contribute indirectly to disease pathology. Despite significant progress, seven OFD syndromes are still without a clear genetic aetiology and at least three unclassified forms need further clinical and molecular characterization.

The Integrator complex (INT), which is unique to metazoans, is composed of at least 14 subunits, and it associates with RNA polymerase II (RNAPII) where it modulates transcription^25, 26, 27, 28, 29^. Critical among its protein components is Integrator subunit 11 (INTS11)^30^, which is a member of the metallo-β-lactamase/β-CASP family of RNA endonucleases^31^ and is a paralog of the cleavage and polyadenylation specificity factor of 73 kDa (CPSF73), which cleaves pre-mRNA^32^. INTS11 has been found to associate with INTS4 and INTS9 to form what is called the Integrator Cleavage Module (CM), which is thought to be the key enzymatic factor within the complex^33^. The Integrator CM, through the activity of INTS11, has been shown to be essential to cleave the 3’ ends of a variety of noncoding RNA and nascent mRNA of protein-coding genes with paused RNAPII. This catalytic activity thus serves as a key functional component of Integrator-mediated transcriptional repression and for the biogenesis of noncoding RNA^34, 35, 36, 37, 38^.

The potential connection between Integrator function and ciliogenesis was uncovered initially in *Drosophila*, where depletion of nearly any Integrator subunit disrupts proper centriole localization, an early event in ciliogenesis^39, 40^. Further evidence was attained using human retinal pigment epithelial (RPE) cells, where depletion of multiple Integrator subunits can lead to inhibition of primary cilium biogenesis^41^. More recently, an unbiased CRISPR screen for factors required for ciliogenesis also identified several Integrator subunits as essential to form primary cilia^42^. Collectively, these data imply that the overall function of the Integrator complex, and not any specific subunit, is required for proper cilia formation. Given its role in modulating RNAPII behavior at diverse gene types, the precise function for Integrator in ciliogenesis is most likely at the level of gene expression. However, the ciliary genes subject to Integrator regulation are currently unknown, and no ciliopathies have been found associated with Integrator subunit mutations.

Here, we used a combination of homozygosity mapping and deep sequencing of patient samples to identify two distinct recessive germline mutations in the *INTS13* (MIM615079) gene causing a variable form of OFD syndrome type 2 (OFD2), also known as Mohr Syndrome (MIM252100)^43^. Using multiple molecular and biochemical approaches, we demonstrate that the C-terminus of INTS13 interacts with the Integrator CM and that mutations observed in OFD2 patients specifically disrupt this interaction. We also find that human cells depleted of INTS13 not only have reduced primary cilia, but also exhibit broad transcriptional dysregulation that impacts numerous known genes required for proper ciliogenesis. These observations were validated *in vivo*, where morpholino-mediated depletion of *ints13* in *Xenopus laevis* leads to profound underdevelopment, with a specific reduction in the formation and beating of multiciliated cells. Taken together, these results reveal a key role for Integrator in human ciliogenesis and identify a vital interaction between two Integrator subcomplexes.

## RESULTS

### Characterization of 4 patients with recessive loss-of-function *INTS13* mutations

In Family 1, two affected sisters (individuals II.4 and II.5 in Figure 1a) born to a consanguineous Jordanian couple presented with a congenital disorder consistent with OFD2 (MIM252100). The two sisters displayed similar phenotypes encompassing craniofacial dysmorphisms, including: a bilateral cleft lip (CL), microcephaly, hypertelorism, epicanthal fold, broad nasal root with depressed nasal tip, large and low set ears, long philtrum and thick septum. In addition to bilateral CL, their oral cavity had omega shape epiglottis, underdeveloped epiglottic folds and hypertrophied false vocal cords. Dental abnormalities were also observed, such as central median upper incisor in II.4 and single central upper incisor in II.5. Supernumerary teeth were observed in II.4. Both sisters have clinodactyly of the 4^th^ and 5^th^ fingers and toes, and pre-axial polydactyly was observed in one hand of II.4. Single transverse palmar creases on both hands and rough hair were also observed in both sisters. Audiologic examination of the two sibs revealed the presence of tympanic membrane retraction and calcification (myringosclerosis). Ophthalmic examination revealed dilated retinal vessels and a crowded optic disk. Patients were reported to have chronic urinary tract infection and recurrent respiratory tract infections with chronic cough. Supravalvular pulmonary stenosis and partial anomalous pulmonary venous return were reported in patient II.5 soon after birth. Taken together, both patients share the cardinal features of an Oral-Facial-Digital Syndrome and exhibit a wide range of clinical manifestations. The clinical findings of these patients are illustrated in Supplemental Figure 1a-m, and a summary is provided in Table 1. Inter- and intra-familial variations have been reported in OFD and were clearly observed between the two sisters. Many clinical features are unique to this family and overlap with OFD2 or may be a new form of OFD, with a distinct genetic aetiology.

**Fig 1.**
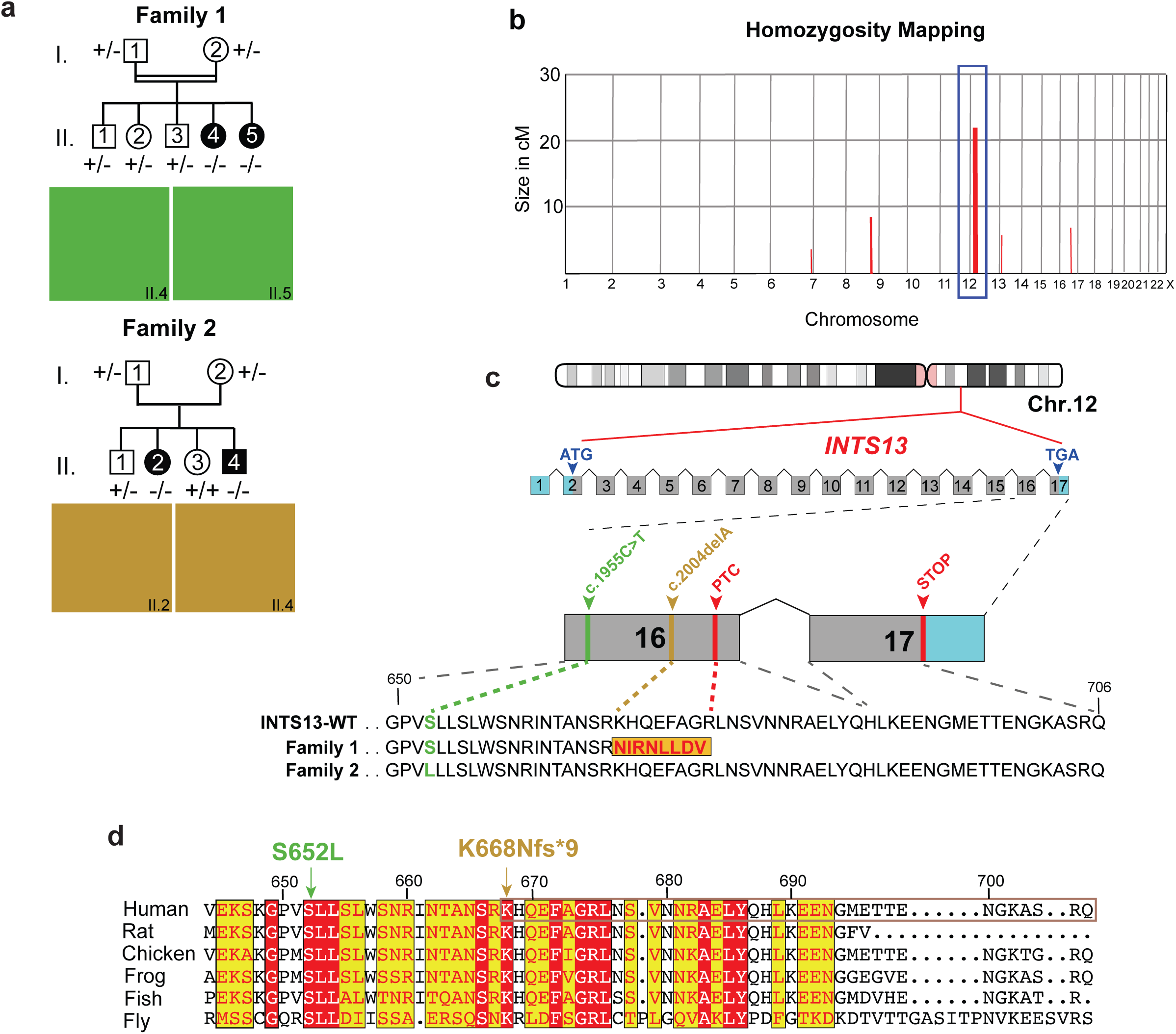
Identification of two families segregating recessive mutations in *INTS13*. **a** Pedigree of Family 1 (top): affected individuals (II.4 and II.5) were born to first-cousin parents (I.1 and I.2) with three unaffected children (II.1-II.3). Double lines indicate a consanguineous marriage. Pedigree of Family 2 (bottom): affected individuals (II.2 and II.4) were born to parents with two unaffected children (II.1 and II.3). **b** Homozygosity mapping of Family 1 delineated five candidate loci totaling 46 cM on chromosomes 6, 8, 12, 13, and 16. **c** Schematic representation of human *INTS13* located on chromosome 12 (ch12p13.2-p11.22) consisting of 17 exons. Locus capture followed by massive parallel sequencing identified a homozygous frameshift mutation as a disease-causing mutation in Family 1. The c.2004delA (p.K668Nfs*9) mutation is caused by a single base pair deletion in exon 16 (marked in brown), resulting in a frame shift and premature termination codon (PTC) which alters nine amino acids (marked in red) and deletes the last 31. In Family 2, the c.1955C>T (S652L) mutation is caused by a single base pair substitution in exon 16 (marked in green). **d** Amino acid alignment of the C-terminus of the INTS13 protein shows that both mutations occur at highly conserved residues. Red and yellow shading indicates regions of conservation.

To identify the genetic origin of this developmental disorder, the parents, both affected siblings, and healthy siblings were screened on a 610k Illumina SNP array. A subsequent homozygosity identical-by-descent (IBD) mapping revealed five homozygous candidate regions that were unique to the proposita, with a total size of 45.76 Mb on chromosomes 6, 8, 12, 13 and 16 (Figure 1b). We next employed targeted genomic loci capture followed by massive parallel sequencing to screen 395 candidate genes in these loci^44^. Out of 894 mismatches, one homozygous missense mutation (c.6585A>T) in *SACS* (MIM604490) and a frameshift (c.2004delA) mutation in *INTS13* (MIM615079) were found in the two affected sisters. The mutation in *SACS* was ruled out as the likely cause due to the gene’s known role in causing spastic ataxia (MIM 604490), which did not align with the patients’ phenotypes^45, 46^. The single base pair (bp) deletion in exon 16 of *INTS13* was never observed in GnomAD and is predicted to be damaging according to SIFT or MutationTaster. It alters the reading frame, leading to a premature termination codon (PTC) at p.K668Nfs*9 and the deletion of the remaining 31 amino acids of the INTS13 protein (Figure 1c). This private allele segregated with the disease in all family members available (Supplemental Figure 1n).

The second family had two affected siblings born to French parents not known to be endogamous. The affected 27-year-old woman (II.2) and her affected 20-year-old brother (II.4) (Family 2, Figure 1a) presented with some overlapping phenotypes, but also distinct features from those reported in the Jordanian patients. Both affected sibs were born at term with low birth parameters for weight, length and occipitofrontal circumference (OFC). II.4 presented with hypospadias. Both remained short and became overweight from the age of 3 years, while OFC was in the low normal range at this age. II.2 walked unaided before 18 months and II.4 walked at 22 months. For both, language was delayed and remained limited to a few spoken words, although they could understand simple conversational topics. II.2 was treated for early-onset puberty for 18 months starting at 8 years. Both presented with short extremities with brachydactyly of fingers and toes, hyperlordosis when standing, flessum of the knees and pes valgus. Facial features include upslanting palpebral fissures, macrostomia, gum hypertrophy and a bulbous nasal tip. They wear glasses for correction of hyperopia and astigmatism. At 20 years, regression with less social interactions, vocal stereotypies and tip toe walking were noted in II.4, while his brain CT-scan showed some cerebellar atrophy and calcification of the cerebellum tent. II.2 has remained stable with no regression observed thus far. The clinical findings of these patients are illustrated in Supplemental Figure 1o-w, and a summary is provided in Table 1.

Agilent 60k Array CGH showed no disequilibrium in either sibling. Exome capture was performed followed by sequencing of both affected siblings and their healthy parents. Each sample had an overall mean depth of coverage greater than 90X and more than 97% of the exome covered at least 15X. Filtering for germline compound heterozygous variants, X-linked or shared heterozygous de novo variants did not provide any candidate; however, a single shared homozygous variant in *INTS13* (NM_018164.2: c.1955C>T) emerged as the only candidate mutation (Figure 1c). This mutation converts Serine652 to a Leucine and is located 16 amino acids upstream of where the frameshift mutation was observed in Family 1.

*INTS13* mRNA consists of 17 exons (Figure 1c). Its encoded protein is 706 amino acid residues long and was initially termed Mat89Bb (Maternal transcript 89Bb) due to its maternal expression in *Drosophila* embryos^47, 48^. Mat89b was later changed to Asunder/ASUN given its mutant phenotype of altered association of the nuclei with asters and disrupted spermatogenesis^49^. Importantly, ASUN was later shown to be a component of the Integrator complex, although little is known of its distinct function in that context^50^. There is an overall high degree of sequence similarity within the C-termini of INTS13 proteins found in multiple organisms, and both mutated residues are highly conserved between vertebrate and invertebrate species (Figure 1d), suggesting that alteration of these amino acids may lead to a pathogenic function of the INTS13 protein.

### *INTS13* mutations affect endogenous INTS13 protein levels in patient cells

To investigate the impact of *INTS13* mutations on its expression, we isolated primary cutaneous fibroblasts from affected patients in both families as well as fibroblasts from healthy individuals. We first compared mRNA expression levels by RT-qPCR of *INTS13* in Family 1 cells (F1) containing the INTS13 truncation mutation. We found the level of *INTS13* transcripts to be significantly down-regulated (Figure 2a), which we hypothesized may be due to the presence of the premature termination codon triggering nonsense-mediated decay (NMD). Therefore, we isolated total RNA from patient cells that were either mock-treated or treated with cycloheximide, a translational inhibitor that blocks NMD, and RT-qPCR was repeated. The results showed that the level of mutant mRNA was increased following cycloheximide treatment, suggesting that the mutant *INTS13* mRNA was indeed targeted for NMD (Figure 2b). To assess the levels of INTS13 protein in F1 cells, we raised two custom antibodies against antigens from the central portion of INTS13 (α-M) and the C-terminal region (α-C), which is downstream of the truncation. These antibodies were affinity-purified and used to probe cell lysates from patient fibroblasts using Western blot analysis. We observed that the C-terminal antibody could detect endogenous INTS13 protein in control cells, but not in F1 patient fibroblasts, likely due to the loss of the epitope in the truncated mutant (Figure 2c). Consistent with this notion, we could detect low levels of truncated INTS13 in F1 patient fibroblasts using the α-M antibody (Figure 2c). Overall, these results reveal that the *INTS13* mutation observed in Family 1 creates an mRNA subject to NMD that is translated at low levels, thereby producing a truncated INTS13 protein.

**Fig 2.**
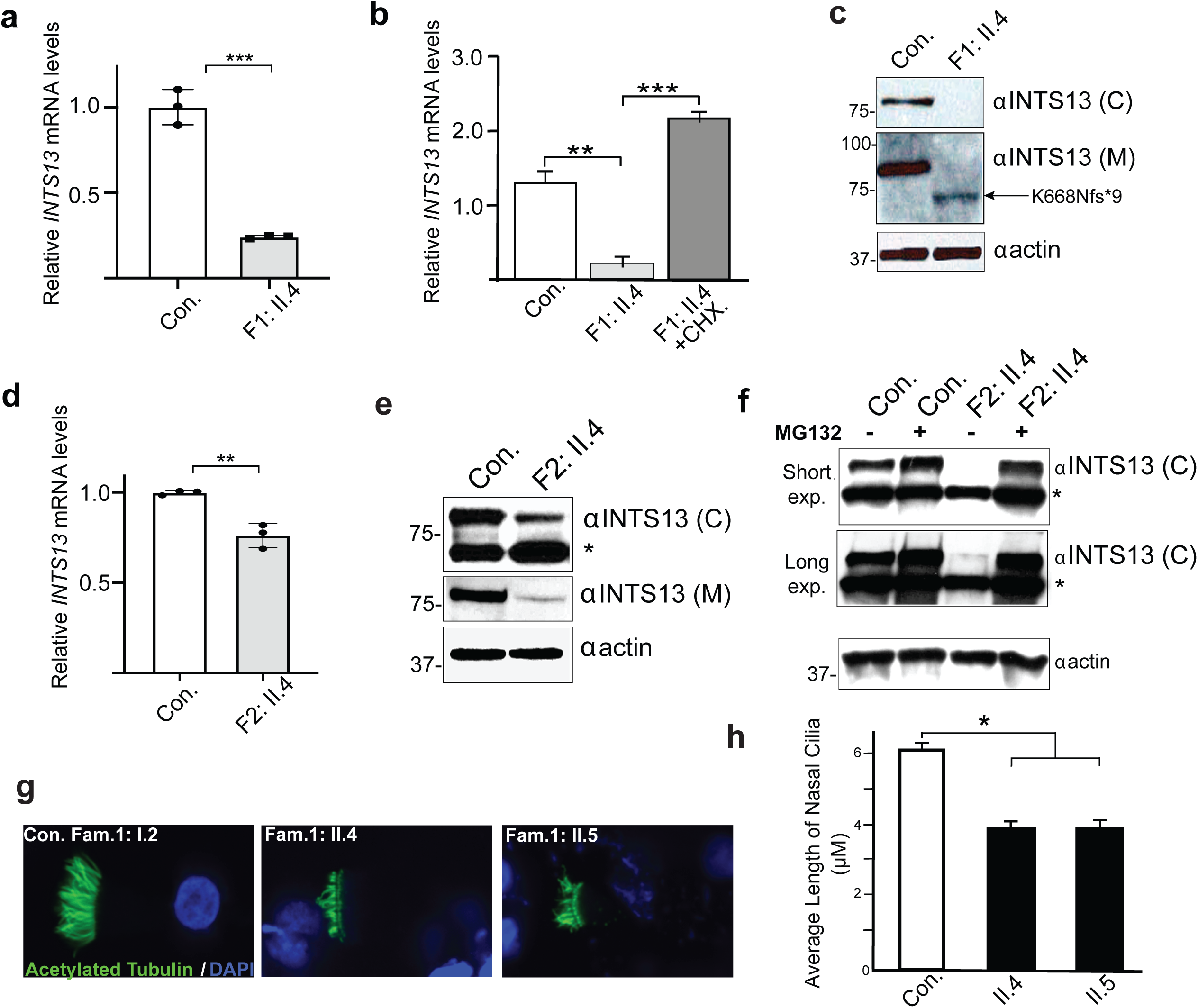
Characterization of Loss-of-function INTS13 mutations in patient cells. **a** RT-qPCR shows that endogenous *INTS13* transcript levels were significantly reduced relative to control cells in primary dermal fibroblasts of affected individual II.4 from Family 1. Data are mean +/-SD, n = 3. Statistical significance was calculated using a two-tailed unpaired t-test. **b** Primary dermal fibroblasts from individual II.4 from Family 1 were treated with 100 g/ml cycloheximide (CHX). Following treatment, RT-qPCR was done which shows that the *INTS13* mRNA level was increased in treated cells. Data are mean +/-SEM, n = 3. Statistical significance was calculated using a one-tailed Student’s t-test. **c** Western blot analysis of primary dermal fibroblasts of affected individual II.4 from Family 1 using two polyclonal antibodies designed against distinct regions of INTS13. No signal was detected in patient cells with the antibody directed against the C-terminus of INTS13 (C). A truncated protein was detected in patient cells by the antibody directed against the center of the protein (M). Actin was used as a loading control. **d** RT-qPCR shows that *INTS13* transcript levels were significantly reduced in primary dermal fibroblast cells of affected individual II.4 from Family 2 relative to control cells. Data are mean +/-SD, n = 3. Statistical significance was calculated using a two-tailed unpaired t-test. **e** Western blot analysis of primary dermal fibroblasts of affected individual II.4 from Family 2 shows reduced levels of INTS13 protein compared to control cells. **f** Western blot analysis of primary dermal fibroblasts from individual II.4 from Family 2 treated with MG132. Treated cells show a level of INTS13 protein comparable to control cells. * indicates a non-specific signal. **g** Multiciliated airway cells were obtained from nasal biopsies of three members of Family 1: the unaffected carrier mother (Con.) and the two affected children (II.4 and II.5). Probing for acetylated α-tubulin shows significantly shorter, less dense, and disorganized cilia in the affected individuals compared to the mother’s cells. **h** Bar graph representing the average length of nasal cilia. Data are mean +/-SEM, n = 200 cells for each individual. Statistical significance was calculated using a one-tailed t-test. *, **, ***, **** correspond to p-values < 0.05, 0.01, 0.001, and 0.0001, respectively.

We next analyzed the expression of INTS13 in fibroblasts isolated from a Family 2 (F2) patient harboring the homozygous p.S652L mutation. We observed a slight but significant reduction in the level of endogenous *INTS13* mRNA in these mutant cells (Figure 2d) and a pronounced reduction in the level of INTS13 protein as assessed using either antibody (Figure 2e). We posited that the basis for the reduction in INTS13 protein, beyond the minor reduction in mRNA, may be due to increased protein degradation of the INTS13 F2 mutant. To test this, primary fibroblasts were treated with MG132 to inhibit proteasome activity, and we observed an increased level of INTS13 protein comparable to control cells by Western blot analysis (Figure 2f). This suggests that the INTS13 mutation observed in Family 2 generates a protein which may be unstable and targeted for degradation via the proteasome.

The epithelia lining the human respiratory system are covered with multiple cell types dominated by ciliated cells. These ciliated cells have hundreds of motile cilia to generate fluid flow that keeps mucus moving to protect the airway from inhaled irritants, pathogens and foreign particles. Abnormalities of mucociliary clearance caused by either aberrant ciliated cells or cilia dysfunction often result in recurrent respiratory tract infections. Reduction or impairment of mucociliary clearance in humans results in an array of pathologies such as primary ciliary dyskinesia, chronic obstructive pulmonary disease and asthma ^51, 52, 53, 54, 55^. In this study, both probands of Family 1 suffered from recurrent respiratory tract infection and chronic cough. Therefore, we obtained respiratory epithelial cells from the two patients and the mother to assess cilia formation and performance. Immunofluorescence staining was performed using anti-acetylated α-Tubulin antibody to visualize cilia. The average length of nasal cilia was then scored. The results showed that cilia were shorter, fewer in number and occasionally disorganized compared to those of the heterozygous but unaffected mother, which may explain the recurrent respiratory tract infections in the two affected subjects (Figure 2g-h). Importantly, these results also indicate that ciliogenesis was indeed disrupted in patient cells.

### OFD2 patient mutations within the INTS13 C-terminus disrupt association with the Integrator Cleavage Module

Recently, the structure of INTS13 associated with INTS14 has been elucidated. However, the INTS13 C-terminus was unable to be resolved in that structure, thereby obscuring our ability to ascertain the impact of OFD2 mutations^56^. The INTS13/14 heterodimer was shown to interact with INTS10 as well as the Integrator CM, the latter of which was found to be mediated independently through the INTS13 C-terminus^56^. With this information in hand, and our previous success mapping Integrator interactions using yeast two-hybrid analysis^33, 50, 57, 58^, we fused INTS13 to the Gal4 DNA binding domain and screened for interactions with the other Integrator subunits that were each expressed individually as Gal4 activation domain fusions. This initial approach failed to detect interaction between INTS13 and other members of the complex (Figure 3a, left panels). Given the structural information for INTS13, the most likely explanation for the inability to detect binding partners is that INTS13 may interact with a domain created by an interface between multiple Integrator subunits. We previously demonstrated this to be the case for the INTS4/9/11 Cleavage Module where INTS4 interaction with INTS9 or INTS11 was only detected with yeast two-hybrid if all three subunits were co-expressed^33^. We therefore decided to focus on whether INTS13 could interact with members of the cleavage module and thus repeated the screen with INTS9 and INTS11 expressed simultaneously in *trans*. Using this approach, we could detect a robust interaction between INTS13 and INTS4 only when INTS9 and INTS11 were co-expressed in the same yeast strain (Figure 3a, right panels and Supplemental Figure 2a). To confirm this interaction, we altered the position of each subunit in the screen and could detect an interaction between INTS9 and INTS13 only when both INTS4 and INTS11 were simultaneously expressed in *trans* (Supplemental Figure 2b). These data extend on recent structural and biochemical analyses to indicate that INTS13 can, in the absence of INTS10 and INTS14, associate with subunits of the Integrator CM.

**Fig 3.**
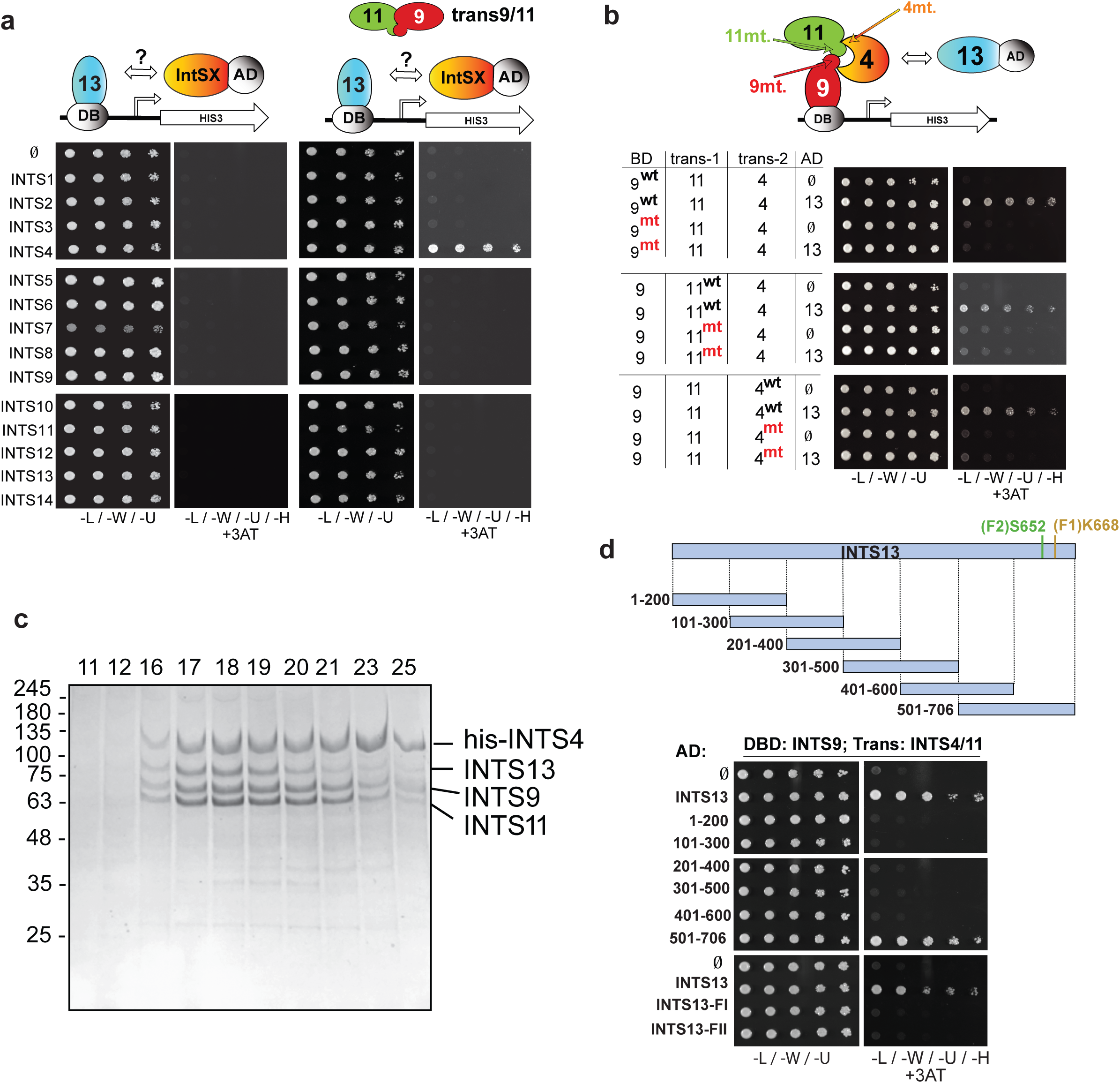
INTS13 associates with the INTS4/9/11 heterotrimer. **a** Yeast two hybrid screening for interactors of INTS13. Haploid strains of yeast expressing each individual Integrator subunit fused to the Gal4 activating domain (AD) were mated with strains expressing INTS13 fused to the DNA binding domain (BD). The first and third panels show mated yeast grown on selective media as a positive control (media lacking leucine, tryptophan, and uracil). The remaining panels are of yeast plated on restrictive media additionally lacking histidine to test for protein-protein interactions and with 3AT (3-amino-1,2,4-triazole) to test the strength of the interaction. No interaction was observed between INTS13 and any other individual subunit (second panel). When INTS9 and INTS11 were expressed in *trans* with INTS13, there was a positive interaction with INTS4 (fourth panel). **b** Point mutants of INTS4, INTS9, and INTS11 known to disrupt their interactions were used in yeast two hybrid to demonstrate that the formation of the INTS4/9/11 heterotrimer is necessary for INTS13 association. General locations of the mutations are indicated with arrows on the schematic. INTS13 is fused to the activating domain in the rest of the yeast two hybrid assays shown in this study. **c** SDS PAGE gel of fractions from gel filtration column of co-expressed full-length INTS4, INTS9, INTS11 and INTS13. The sample was first purified by nickel affinity chromatography. INTS4 carried an N-terminal His-tag, while the other three proteins were untagged. The positions of the fractions are indicated at the top of the gel. **d** Consecutive 200-amino acid truncations of INTS13 were tested for interaction with INTS4/9/11. In modified yeast two-hybrid assays, the only positive interactions seen are full length INTS13 and the last ∼200 residues, 501-706. The patient mutations are marked on the schematic and occur in this region. Loss of interaction with INTS4/9/11 was observed using the mutant versions of INTS13.

The binding interfaces of the Integrator CM have been previously well-characterized using both biochemical and structural approaches^33, 57^. Based upon these prior analyses, amino acid mutations within each of the three CM subunits are known to disrupt formation of the heterotrimer. Therefore, we repeated our modified two-hybrid assay with each of these mutants tested individually and discovered that any disruption of the INTS4/9/11 CM also results in loss of INTS13 interaction (Figure 3b). Further, in order to confirm the interaction between these four subunits using an independent approach, we co-infected High Five cells with baculoviruses individually expressing His-tagged-INTS4, or untagged INTS9, INTS11, and INTS13. The complex was purified from cell lysate using nickel affinity resin and subjected to gel filtration analysis. We observed that all four Integrator subunits remained intact as a complex and could be detected in the same fractions at near stoichiometric levels (Figure 3c). Collectively, these results indicate that INTS13 interacts with the three components of the Integrator CM only once the module is fully assembled.

We next wanted to determine which region of INTS13 is responsible for this interaction and whether OFD2 mutations may impact it. We tested a series of INTS13 fragments in modified yeast two-hybrid assays to determine whether any of these regions were sufficient to mediate interaction with INTS4/9/11 (Figure 3d schematic). The INTS13 C-terminal region, composed of amino acids 501-706, was as effective as the full-length INTS13 to support growth on selective media (Figure 3d). We further dissected the INTS13 C-terminus through a series of smaller deletion mutants, which ultimately defined that the last 130 residues, amino acids 577-706, was sufficient to bind to the Integrator CM (Supplemental Figure 2c). This result is consistent with biochemical approaches recently described where a GST fusion protein containing the C-terminus of INTS13 can pull down subunits of the Integrator CM^56^. Because both mutations observed in Family 1 (p.K668Nfs*9) and Family 2 (p.S652L) are within the region of INTS13 found to be sufficient to bind INTS4/9/11, we tested whether these mutations could disrupt binding. Indeed, we found that either mutation in the context of the full-length INTS13 protein resulted in a loss of interaction between INTS13 and INTS4/9/11, indicating that this interaction may be specifically disrupted in the four affected patients (Figure 3d).

### The INTS13 C-terminus connects an INTS10/13/14 submodule to the Integrator complex

We next sought to determine if the C-terminal INTS13 mutations have an impact on the assembly of the entire Integrator complex. To do so, we created individual 293T cell lines harboring three different versions of the *INTS13* cDNA: wild type (wt), Family 1 mutant (F1), or Family 2 mutant (F2). Each of these lines expresses exogenous INTS13 with an N-terminal FLAG tag under the control of the doxycycline-inducible promoter. After 48 hours of doxycycline treatment, nuclear fractions from each of the cell lines as well as an untransfected naïve line serving as a negative control were extracted. Purified Integrator complexes associated with each INTS13 protein, and the degree of co-associated INT subunits was assessed by LC/MS and Western blot analysis. We observed that both patient-derived INTS13 mutants not only disrupted association with INTS4/9/11, confirming our yeast two-hybrid experiments, but also resulted in reduced binding to nearly all other members of the complex (Figure 4a). The exception to this observation was the binding of INTS14 and to a lesser extent INTS10, which were not as dramatically impacted by the mutations (Figure 4a).

**Fig 4.**
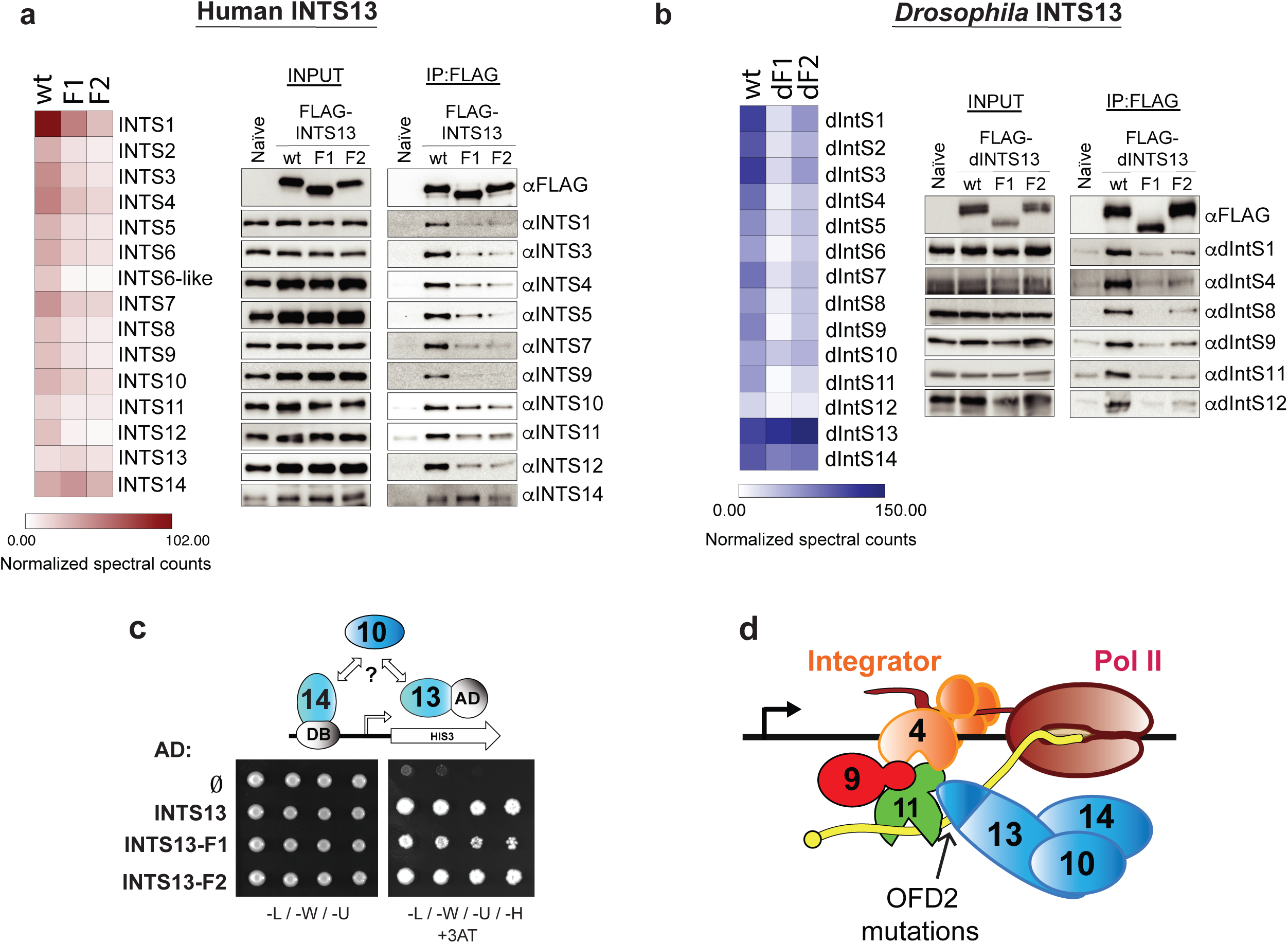
OFD2 mutations lead to conserved disruption of interactions between the INTS10/13/14 module and the remainder of the Integrator complex. **a** Immunoprecipitation of FLAG-INTS13 from nuclear extract derived from HEK-293T cells stably expressing FLAG epitope tagged human INTS13-wild type (wt), Family 1 INTS13 mutant (F1), or Family 2 INTS13 mutant (F2). Nuclear extract derived from naïve HEK-293T cells not expressing any FLAG epitope was used as a control. Left panel: IP-LC/MS was done for each sample, and quantitation is shown as a heat map. The color scale shown denotes the normalized spectral counts for each protein (see methods). Right panel: Western blot analysis of either nuclear extract input or anti-FLAG immunoprecipitation using human Integrator antibodies. **b** Results shown are from parallel experiments conducted in a manner identical with panel (a) with the exception that nuclear extract was derived from *Drosophila* S2 cells expressing FLAG-dINTS13-wt or FLAG-tagged dINTS13 proteins where patient mutations were introduced into homologous regions of the *Drosophila INTS13* cDNA. **c** Modified yeast two hybrid analysis where human INTS14 is expressed as a Gal4-DNA binding domain fusion, human INTS13 wt or mutant is expressed as a Gal4-Activation domain fusion, and human INTS10 is expressed in *trans*. Dilutions of yeast are plated on both permissive (left) and restrictive (right) media. Empty activation domain vector is used as a negative control. **d** Schematic of the Integrator CM associating with the remainder of Integrator subunits that is bridged to the INTS10/13/14 module through the C-terminus of INTS13, which is disrupted by OFD2 mutations.

We have shown previously that Integrator constituency and function is highly conserved in *Drosophila*^50^. Therefore, we wanted to test if patient mutations have the same disruptive effect in the orthologous fly complex. Supporting this approach, the C-terminal region of INTS13 is conserved in the *Drosophila* IntS13 orthologue (Asunder or dIntS13) and so we introduced the analogous patient mutations into dIntS13 (Supplemental Figure 3a). We then created individual S2 cell lines expressing copper-inducible FLAG-dIntS13-wild type (wt), Family 1 mutant (dF1), or Family 2 mutant (dF2). We generated nuclear extract from these cell lines as well as a naïve S2 control and purified *Drosophila* Integrator complexes using anti-FLAG affinity resin. Similar to the human Integrator purifications, we used both LC/MS and Western blot analysis to assess complex integrity. In accordance with the human data, we found that patient mutation ‘equivalents’ within *Drosophila* IntS13 reduced association with most other Integrator subunits (Figure 4b). We also observed that association of dIntS10 and dIntS14 were only marginally affected by the patient mutations.

These results confirm that OFD2-causing mutations disrupt INTS13 assembly into Integrator, but also suggest that INTS13 mutants do not affect INTS14 and possibly INTS10 binding. Therefore, we wanted to determine whether we could detect an interaction of INTS13 with INTS10 and INTS14 using our modified two-hybrid assay and if INTS13 mutations affected it. We initially expressed INTS14 fused to the Gal4 DNA binding domain and screened for interactions individually with each of the other Integrator subunits fused to the activating domain. This approach did not detect any pairwise interactions (Supplemental Figure 3b, left panel). Therefore, we expressed each Integrator subunit individually *in trans* with INTS14 fused to the DNA binding domain and INTS13 fused to the activating domain, and we found that the additional presence of INTS10 can allow for an interaction between INTS14 and INTS13, supporting the model that these three subunits form a module as has been recently shown biochemically^56^ (Supplemental Figure 3b, right panel). Finally, we tested whether our patient-derived mutations within INTS13 affect the interaction of INTS10/13/14 and observed that both INTS13 mutants could interact with INTS10/14 comparable to the wild-type INTS13 (Figure 4c). Overall, the results presented here provide a molecular basis for OFD2 where INTS13 mutations disrupt interaction between the INTS10/13/14 submodule and the INTS4/9/11 CM, along with the rest of the Integrator complex (Figure 4d).

### Reduced INTS13 expression in RPE cells leads to loss of ciliogenesis and broad disruption of transcriptional regulation of ciliary genes

Results presented thus far reveal that OFD2 patient cells exhibit disrupted ciliogenesis and that these same mutations also disrupt the INTS13 module’s interaction with the CM. We next wanted to determine whether reduced expression of INTS13 in cultured cells was sufficient to inhibit ciliogenesis. We utilized human retinal pigment epithelial (RPE) cells as a model, because these cells exhibit clearly visible primary cilium upon serum starvation. We treated RPE cells with either control siRNA or one of two distinct siRNA targeting *INTS13* and discovered that either siRNA could effectively deplete endogenous INTS13 (Figure 5a). To visualize primary cilia, we probed fixed cells using antibodies raised to ADP-ribosylation factor-like protein 13B (ARL13B)^59^, which stains the ciliary axoneme, and γ-tubulin, which stains the basal body^39^. We observed that the basal body and axoneme components were still present and appropriately juxtaposed in INTS13-depleted cells but were clearly not utilized to form a normal primary cilium (Figure 5b). We found that depletion of INTS13 with either siRNA reduced the number of properly assembled primary cilium by ∼50% relative to control siRNA (Figure 5c).

**Fig 5.**
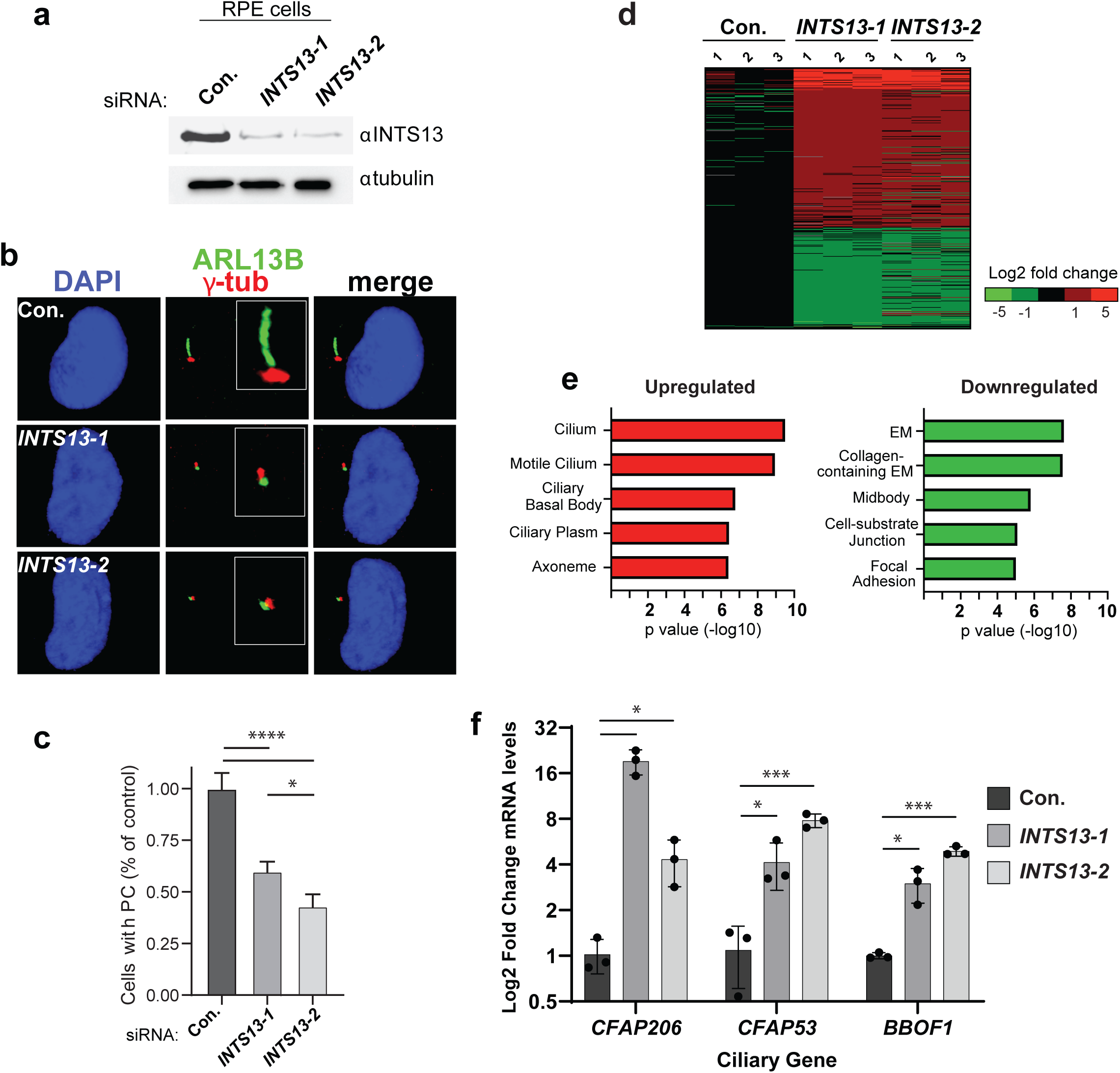
INTS13 depletion leads to a loss of primary cilia and broad transcriptional perturbation of ciliary genes. **a** Western blot of whole cell lysates from RPE cells treated with non-targeting control siRNA (Con.) or either of two siRNA targeting *INTS13* (*INTS13-1, INTS13-2*). Lysates were probed for either INTS13 or tubulin as a loading control. **b** Immunofluorescence imaging of RPE cells treated with siRNAs as in panel (a) to visualize primary cilia (PC). DAPI staining in blue indicates the nuclei, γ-tubulin staining is shown in red to visualize the basal body, and ARL13B staining is shown in green to visualize the ciliary axoneme. Representative images show disrupted primary cilia that occurred when INTS13 was targeted by RNAi. **c** Images from panel (b) were quantified for each condition and graphed as a percentage of the number of control-treated cells with PC (data are mean +/-SD, con. n = 3785, 13-1 n = 2969, 13-2 n = 2675. Raw calculations for each condition were divided by the control mean to adjust control to 1). Statistical significance was calculated using a one-way ANOVA with multiple comparisons. **d** RNA sequencing was done on siRNA treated RPE cells in triplicate. Genes with significant changes in expression are shown by color indicating the log2 fold change. **e** Results from panel (d) were used for gene ontology analysis: the top five significant results are shown for upregulated and downregulated genes. All five GO terms for upregulated genes are related to cilia. **f** Quantitative RT-PCR validating three ciliary genes with expression changes identified using RNA-seq. Results are quantified from independent biological replicates (n = 3, data are mean +/-SD). Statistical significance was calculated using multiple Student t-tests with a Bonferroni correction. *, **, ***, **** correspond to p-values < 0.05, 0.01, 0.001, and 0.0001, respectively.

To understand the underlying changes in gene expression that occurred after *INTS13* depletion, we subjected control siRNA-treated and each of the *INTS13* siRNA-treated RPE cells to RNA-seq analysis. We conducted RNA-seq using three biological replicates from cells treated with each siRNA and used DESeq to determine differential expression (Supplemental Table 1). Taking advantage of two distinct *INTS13* siRNAs, we only considered genes observed to be significantly upregulated/downregulated (>2-fold, p<0.0001) in both knockdown conditions. Using these filters, we identified 254 genes that were upregulated and 156 genes that were downregulated upon *INTS13* knockdown (Figure 5d). Gene ontology analysis revealed a striking enrichment in genes involved in cilium biogenesis uniquely within the upregulated gene set, suggesting that knockdown of *INTS13* leads to de-repression of a collection of genes involved in cilium biogenesis (Figure 5e). We validated several of these gene changes using RT-qPCR including well-characterized ciliary genes such as *BBOF1, CFAP206*, and *CFAP53* (Figure 5f). Altogether, these results indicate that depletion of INTS13 in human cells leads to disrupted ciliogenesis that is likely caused by broad transcriptional deregulation of genes involved in primary cilium biogenesis.

### Knockdown of *ints13* causes developmental defects and ciliary phenotypes in *Xenopus*

To examine the role of INTS13 *in vivo*, we first examined its expression in *Xenopus laevis* embryos using whole mount in situ hybridization (WISH). *ints13* showed both maternal and zygotic contributions with relatively ubiquitous expression. At late gastrula (stage 12), the expression of *ints13* was enriched in ectodermal lineages, with no apparent expression in the endoderm. At neurula (stage 17), its expression was enriched in the neural tube and neural folds (Supplemental Figure 5a). From tadpole stages onwards, *ints13* transcripts were abundantly observed in ciliated tissues and organs including the brain, branchial arches, pronephros, eye and otic vesicles. Remarkably *ints13* was found to follow a punctate pattern of expression in the skin, which coincided with that of multiciliated cells (MCCs). This was validated by whole mount antibody staining, using the α-M antibody highlighting that endogenous INTS13 was enriched in epidermal cells endowed with multicilia (Supplemental Figure 5b).

In order to gain insight into the precise function of ints13 during *Xenopus* embryonic development, we performed loss-of-function analysis using a morpholino oligonucleotide (MO) targeted against the *ints13* start codon (IntS13-MO). This IntS13-MO was previously established to be specific^47^, and we confirmed the translational block efficiency achieved by this MO by demonstrating that *ints13* morphants were nearly devoid of endogenous ints13 protein (Figure 6d). The Ints13-MO resulted in developmental delay, a small head, shortened body axis, bent tail, cardiac oedema, polyploidy and early developmental arrest in *Xenopus* morphants (Figure 6a). In addition to the previously described gastrulation defects, we also observed prominent deficiencies in ciliogenesis and craniofacial development. Based on the enrichment of ints13 in MCCs, from which protrude hundreds of beating cilia^60, 61^, we examined the possibility that *ints13* may affect multi-cilia formation and function. *In situ* hybridization using a specific MCC marker *ccdc19*^62^ showed that morphants had significantly less *ccdc19* than did control embryos (Figure 6b), which was validated by whole mount antibody staining using the anti-acetylated α-tubulin antibody marking cilia protruding from MCCs (Figure 6c). We noted increased distances between puncta and irregular staining in *ints13*-depleted embryos, suggesting that the epidermis of *ints13-*depleted tadpoles could have fewer differentiated MCCs and/or that these cells harbor fewer cilia. Scanning electron microscopy (SEM) performed on both *ints13* morphants and control embryos revealed that MCCs were defective, with a significantly reduced number of cilia per cell (Figures 6e, f). While the length of cilia seemed to be normal, inconsistency was observed within and between MCCs, suggestive of incomplete penetrance (Figure 6e). Cilia exhibited abnormalities in morphology, protrusion and direction (Figure 6f). Of note, we were unable to detect overt abnormalities in the ultrastructure of the cilium by transmission electron microscopy (TEM) (Figure 6g). These results prompted us to examine the function of the cilia, which normally generate an anterior-to-posterior directional flow over the tadpole’s skin. Using high-resolution imaging, time-lapsed movies revealed that morphants were completely unable to move liquid around their bodies compared to control embryos (Supplemental Movie 1). Moreover, we found that ciliary beating in *ints13*-depleted embryos was slow, erratic and uncoordinated compared to control embryos (Supplemental Movie 2). These data indicate that INTS13 is necessary for MCC differentiation, as well as cilia formation and beating, thus revealing its critical role in ciliogenesis during development.

**Fig 6.**
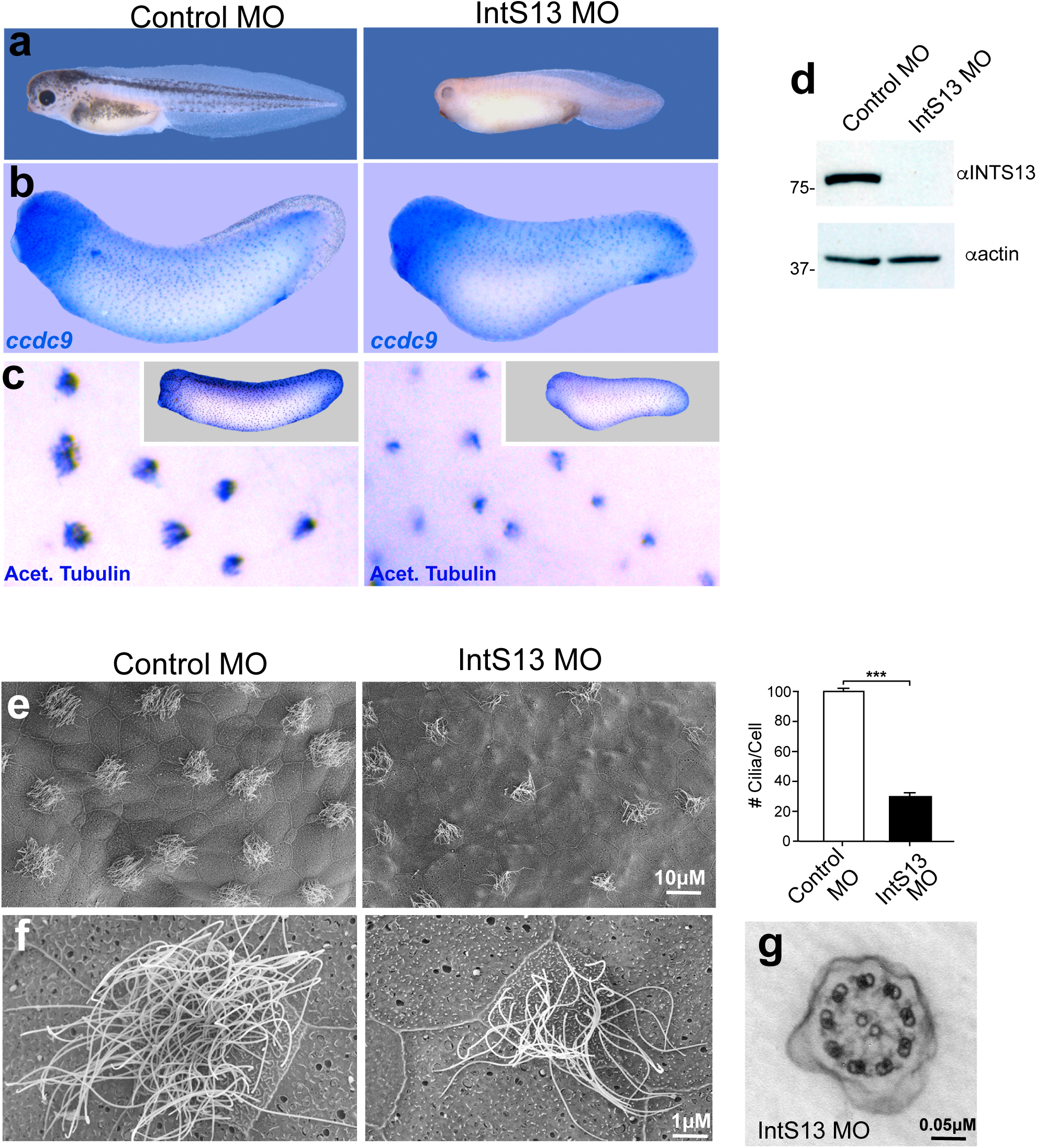
INTS13 is required for embryonic development and ciliogenesis in *Xenopus laevis*. **a** In comparison to control embryos, morpholino (MO)-mediated knock down of INTS13 in *Xenopus* leads to severe developmental defects including a small head, big belly, short and down curved body axes, cardiac edema and massive developmental delay. **b** In situ hybridization using a specific probe for the multi-ciliated cell (MCC) marker *ccdc9* shows a reduction in *ccdc9* expression in MCC of morphants compared to control embryos. **c** Whole mount antibody staining for acetylated α-tubulin shows a punctate pattern similar to *ccdc9*. The puncta have a more condensed and stronger signal in control embryos. A higher magnification view of the MCC shows marked reduction in cilia compared to controls. **d** Western blot showing that the *INTS13* MO was able to deplete INTS13 protein in *Xenopus* embryos. **e** Low magnification SEM imaging of *Xenopus* MCC at embryonic stage 28. MCCs of a control embryo possess multiple, long cilia, whereas the morphant MCCs display fewer cilia per cell which appear to be disorganized. **f** High magnification SEM imaging of *Xenopus* MCC at embryonic stage 28. Ciliary length seems unperturbed in morphant embryos, while the average number of cilia on individual MCCs is clearly reduced by INTS13 knockdown. The average MCC in morphants have less than half of the number of cilia per cell compared to control, as shown in the bar graph. **g** The ultrastructure of the cilia in morphants appears to show a normal 9+2 structure as imaged by TEM. *, **, ***, **** correspond to p-values < 0.05, 0.01, 0.001, and 0.0001, respectively.

## Discussion

Oral facial digital syndromes are an expanding group of genetic conditions characterized by malformations of the face, oral cavity, and digits. These disorders exhibit extensive variability of clinical features with intra-familial and inter-familial variability. Here we describe two homozygous germline mutations in the *INTS13* gene in two independent families, with Family 1 fulfilling clinical criteria for OFD. Through molecular and biochemical approaches, we determined that INTS13 associates with INTS10/14 to form a heterotrimer that interacts with the Integrator CM through the C-terminus of INTS13. The two disease-causing mutations both reduce steady-state levels of endogenous INTS13 protein in patient cells and disrupt INTS13 interaction with the CM, likely causing transcriptional defects in a broad collection of ciliary genes. The binding of INTS10/13/14 to the Integrator CM is clearly vital, as depletion of INTS13 was sufficient to disrupt ciliogenesis in developing *Xenopus* embryos. These results underscore both the importance of Integrator assembly during development and reveal the importance of Integrator transcriptional regulation in ciliogenesis.

INTS13 and INTS14 were not initially described as components of Integrator^25^, but were subsequently discovered through a genome-wide RNAi screen for factors regulating UsnRNA 3’ end formation^50^. Since that initial identification, INTS13 has been shown to play a key role in the differentiation of monocytes^63^. During the course of this manuscript preparation, we became aware of another study in which the structure of INTS13 and INTS14 as a heterodimer was solved^56^. Further, using elegant biochemical approaches, Sabath and colleagues determined that INTS13/14 associates into a heterotrimer with INTS10 and the C-terminus of INTS13 binds to the Integrator CM^56^. These structural and biochemical findings are remarkably concordant with results attained here. Interestingly, it was also discovered that the INTS10/13/14 heterotrimer has an overall architecture related to the Ku70-Ku80 DNA repair complex^64^ and is capable of binding DNA or RNA *in vitro* with the highest affinity for RNA hairpins. Thus, our work here and the structural data in that study establish that Integrator has at least two submodules: the previously defined INTS4/9/11 Cleavage Module containing the RNA endonucleolytic activity and a Nucleic Acid Binding (NAB) module composed of INTS10/13/14. Interaction between these two modules is mediated in part through the C-terminus of INTS13. The structure of the INTS13 C-terminus was not visualized by crystallography, thus obscuring our ability to understand how the germline mutations identified herein disrupt the folding and/or binding of that region^56^. Nevertheless, the collective implication from these two studies is that the Integrator NAB module can bind to RNA hairpins to position the Integrator CM for RNA cleavage and that this process is perturbed in OFD 2/Mohr syndrome patients.

Our results also provide evidence that *INTS13* is a novel ciliopathy gene, and phenotypes associated with *INTS13* mutation are likely due to a dysfunction in Integrator-mediated transcriptional regulation. This finding is consistent with previous studies showing that depletion of multiple Integrator subunits can also lead to disrupted ciliogenesis^41^, and that mutations within INTS1 or INTS8 also create ciliopathy-related symptoms^42, 65^. Herein, we discovered that depletion of INTS13 causes a ∼2-to 10-fold upregulation in a broad collection of ciliary genes in RPE cells, which is consistent with recent work demonstrating that depletion of Integrator subunits in *Drosophila* cells primarily causes an increase in transcription in a subset of genes^37, 38^. We posit that an increase in transcription of a broad collection of ciliary genes is sufficient to disrupt ciliogenesis by potentially overproducing specific components. The concept of dosage imbalance disrupting ciliogenesis has been well documented through reports where gene duplication of centrosome- or cilia-encoding genes can cause ciliopathies^66, 67, 68^. Even small increases in ciliary gene expression can prove disruptive to ciliogenesis, as evidenced by Down Syndrome patients who present ciliopathy symptoms and, due to its location on chromosome 21, overproduce *Pericentrin* a mere ∼1.5-fold^69^. Overall, our results illustrate how sensitive the process of ciliogenesis is to the dosage of components and underscores the need for coordinate transcriptional regulation of genes critical to ciliogenesis. It will be interesting in future studies to determine the mechanism by which Integrator coordinately regulates ciliary genes and how disruption of the NAB and Cleavage Modules gives rise to loss of this transcriptional control.

## METHODS

### Clinical work

The index cases of the Jordanian family were initially diagnosed at the National Center for Diabetes, Endocrinology & Genetics in Jordan by Prof. Hanan Hamamy. Clinical assessments were undertaken at different clinics at the University hospital of Jordan. Saliva samples were collected from the whole family members (I.1, I.2, and II.1 to II.6) and skin biopsies were obtained Family 1 (individuals I.2 and II.4). Family 2 was diagnosed at the Imagine Institute in France by Prof. Jeanne Amiel. Blood and skin biopsy were collected from age-matched healthy individuals and patient II:4 from Family 2. Genomic DNA was extracted using (Puregene DNA Purification Kit, Gentra Systems) following the manufacturer’s instructions. Nasal epithelial cells were also obtained from (Family 1: I.2, II.4 and II.5) by nasal-brush biopsy (Cytobrush Plus, Medscand, Sweden).

### Genomic loci capture

Briefly, custom arrays (Agilent 244K) were designed to target every exonic sequence of the genes present in the five homozygous IBD regions in Family 1 (chr6:167,901,317-170,287,795); (chr8:106,743,988-122,186,235); (chr12:12,129,350-28,091,505); (chr13:21,704,404-26,337,017); (chr16:58,296,974-65,631,053). A DNA library from patient (II:4) was prepared according to the Illumina library generation protocol version 2.3 and hybridized to custom arrays according to the Agilent CGH 244K array protocol, and washed, eluted and amplified. The sample was submitted to one channel of Illumina flow cell and sequenced by Illumina Genome Analyzer (GAII) using standard manufacturer’s protocol. The image data was processed by the provided GA Pipeline (Firecrest version 1.3.4 and Bustard version 1.3.4) and all sequences were aligned to the human reference genome (UCSC, build 18) by Blat-like Fast Accurate Search Tool (BFAST)^70^. Mismatches were further filtered to identify variants seen ten or more times, in which the variant was called as homozygous and did not overlap with a known dbSNP129 entry mismatch. Nonsynonymous mutations were identified with additional SeqWare tools and the ‘‘knownGene’’ gene model from the UCSC hg18. The open-source SeqWare project that provides a LIMS tool for tracking samples (SeqWare LIMS) and a pipeline for sequence analysis (SeqWare Pipeline) was used throughout this work.

For exome sequencing in Family 2, sequences were aligned to the human genome reference sequence (UCSC Genome Browser, GRCh38 build) by Burrows-Wheeler Aligner. Downstream processing was carried out with the Genome Analysis Toolkit (GATK), SAMtools and Picard Tools. Substitution calls were made with GATK Unified Genotyper, whereas indel calls were made with a GATK IndelGenotyperV2. All calls with a read coverage ≤2X and a Phred-scaled single-nucleotide polymorphism (SNP) quality of ≤20 were filtered out. Variants were annotated using an in-house-developed annotation software (PolyWeb) allowing filtering of variants according to relevant genetic models.

### Mutation analysis

Positional candidate genes were obtained from GenBank and Ensembl databases. Segregation analysis was done for candidate mutations by Sanger sequencing with the BigDye Terminator cycle sequencing kit (Applied Biosystems) using primers flanking each mutation. The primer sequences for *INTS13* (NM_018164) and *SACS* (NM_014363) mutations screening are given in Supplemental Table 2.

### Cell culture and transfections

Mouse fibroblast cells and hTERT transformed RPE1 cells were cultured in DMEM and Ham’s F12/DMEM (Invitrogen), respectively. Lymphocytes cells were grown in cell culture medium (RPMI) (Invitrogen). All cultured media were supplemented with 10% Fetal Bovine Serum (FBS) (HyClone), 1X Penicillin / Streptomycin and 2 mM L-glutamine under standard conditions (37°C, 5% CO2). To induce ciliogenesis, cells were grown in serum starvation medium (0.5% FBS) for 72 hours. Transient transfection was done using Lipofectamine 2000 (Life Technologies) according to manufacturer’s recommended protocol.

HEK 293T (ATCC, Manassas VA, CRL-3216) cells were grown at 37°C with 5% CO_2_ in DMEM (Gibco, Waltham MA, #11965-092), supplemented with 10% (v/v) FBS and 1% (v/v) penicillin-streptomycin (Gibco, #15070-063). RPE cells (ATCC, ARPE-19, CRL-2302, lot #70004873) were grown at 37°C with 5% CO_2_ in DMEM/F-12 50/50 (Corning, Manassas VA, 10-092-CV), supplemented with 10% (v/v) FBS and 1% (v/v) penicillin-streptomycin. *Drosophila* S2 cells were grown at 27°C in Sf-900 II SFM (Gibco, #10902-088), supplemented with 1% (v/v) Antibiotic-Antimycotic (Gibco, 15240-062).

293T cells were transfected using CRISPR/Cas9 to make inducible stable lines. 300,000 cells per well were seeded in a 6-well plate. The following day, 2 µl of Lipofectamine 2000 (Invitrogen, Carlsbad CA, 11668-019) was combined with 100 µl of Opti-MEM I (Gibco, 31985-070) per well and incubated at room temperature for 7 minutes. 100 µl of this solution was added to 500 ng of the Cas9 AAVS1 sgRNA plasmid and 500 ng of a pTet-3xFLAG-INTS13 plasmid per well and incubated at room temperature for 20 minutes. The transfection solution was then added to wells containing 1 ml of media. Each insert gene was done in duplicate wells. The next day, duplicate wells were combined, and cells were expanded to a 10-cm dish. 48 hours after transfection, stable cell selection began with 1.5 µg/ml puromycin. Once stable lines were created, cells were maintained at 1 µg/ml puromycin. Human *INTS13* cDNA was mutated by site-directed PCR to create the genes for Family 1 and Family 2. All three versions of the *INTS13* gene, including wild type, were cloned individually into a plasmid containing the tetracycline-inducible Tet-On 3G promotor with a 3x-FLAG tag for insertion into the AAVS1 locus. This was accomplished by co-transfection with the Cas9 plasmid containing the AAVS1 guide RNA (ACCCCACAGTGGGGCCACTA). Protein expression for nuclear extract was induced for 48 hours with 0.4 µg/ml doxycycline for INTS13 wild type and Family 1 and 12 µg/ml for Family 2 in 30 15-cm diameter dishes per line.

Stable lines were also made with S2 cells. *Drosophila* INTS13 cDNA was mutated using site-directed PCR to make orthologous versions of the Family 1 and Family 2 mutations indicated in Supplemental Fig 3. The Family I cDNA also had an early stop codon added to recapitulate the truncation seen in the human gene. These modified genes, and wild type *INTS13*, were cloned into a pMT-3xFLAG-puro plasmid ^37^ following the metallothionein promotor and 3x-FLAG tag. 2×10^6^ cells were plated in Schneider’s media supplemented with 10% FBS in a 6-well plate overnight. 2 μg of plasmid was transfected using Fugene HD (Promega, Madison WI, #E2311). Plasmid DNA was mixed with 8 µL Fugene and 100 µL media and incubated at room temperature for 15 minutes before being added to cells. After 24 hours, 2.5 μg/mL puromycin was added to the media to select and maintain the cell population. Cells were transitioned to SFX media without serum for large scale growth. Protein expression for nuclear extract was induced by adding 500 mM copper sulfate for 48 hours to 1 liter of each cell line grown to approximately 1×10^7^ cells/mL.

### Immunofluorescence staining

For patient cells: cells were seeded on 12 mm cover slips. Cells were grown to around 80% confluency and then incubated with 0.5% serum media for 72 hours to induce cilia formation, washed and kept at 4°C until use. Nasal epithelial cells were suspended in RPMI without supplements to avoid artefacts caused by serum. The cytobrush was moved gently up and down for 5 min to detach ciliary cells from the brush. Samples were then spread onto non-coated glass slides, air dried, and stored at −80°C until use. (See our protocol for immunofluorescence staining, URL)

For RNAi-treated RPE cells (Figure 5b): After RNAi treatment (described in the RNA interference section) and serum starvation of cells grown on cover slips, cells were washed with DPBS and each cover slip was placed in an individual well of a 24-well plate. Cells were fixed to the cover slips by incubating in ice cold methanol for 7-8 minutes and washed thoroughly with PBS. Then samples were blocked for 30 min (all incubation steps done at 37°C with humidity). The blocking buffer used was 1% bovine serum albumin (Sigma-Aldrich, St. Louis MO, A7906-50G) (w/v) in PBS + 0.1% Tween 20 (Fisher Scientific, Fair Lawn NJ, BP337-500). Probing was performed for 2 hours with the following primary antibodies at 1:1000 in blocking buffer: rabbit anti-gamma tubulin (Proteintech, Rosemont IL, 15176-1-AP) and mouse anti-ARL13B (Proteintech, 66739-1-Ig), followed by three washes with PBS for 5 minutes each. Secondary antibody probing was for 1 hour in the dark with the following antibodies at 1:1000 in blocking buffer: goat anti-rabbit Alexa Fluor 594 (Life Technologies, Carlsbad CA, A11037) and goat anti-mouse Alexa Fluor 488 (Life Technologies, A11029), followed by three more washes with PBS. Cover slips were mounted on slides using Fluoroshield Mounting Medium with DAPI (Abcam, Cambridge, UK, ab104139). Confocal imaging was performed on a Leica TCS-SP8 3X system (Leica Microsystems, Wetzlar Germany) equipped with a Leica HC PL APO CS2 63×/1.4 oil immersion objective. A tunable (470-670 nm) pulsed white light laser (Leica WLL) and a UV 405 nm diode laser was used for excitation. Images were processed with the image analysis software, Imaris Pro 9.3.0.

### Reverse transcription (RT-PCR) and Quantitative PCR

Figure 2: Total RNA was isolated from cells or embryos using Trizol (Ambion by Life Technologies, Carlsbad CA, 15596018) followed by extraction using the RNeasy Mini Kit (Qiagen, Hilden Germany). RNA (1 µg) was reverse transcribed using Iscript™ cDNA Synthesis Kit (Bio-transcript lead) and the mis-splicing of the transcript was assessed using pair of primers designed in the exons upstream and downstream of the morpholino binding site. Quantitative real-time PCR was performed with power SYBR green master mix (Applied Biosystems) using a 7300 real-time polymerase chain reaction machine (Applied Biosystems) and normalized to the housekeeping gene *β-actin*. Primer sequences for RT-PCR and quantitative real-time PCR are listed in the Supplemental Table 2.

Figure 5: Total RNA was isolated using Trizol for phenol-chloroform extraction. cDNA was reverse transcribed using Superscript III Reverse Transcriptase (Thermo Scientific, Rockford IL, #18080085) according to the manufacturer’s instructions. Random hexamers were used for cDNA synthesis primers and qRT-PCR was then carried out in triplicate using Bio-Rad iTaq Universal SYBR Green Supermix (Bio-Rad, Hercules CA, #1725120) and measured with the CFX Connect Real-Time System (Bio-Rad). Data was analyzed using the ΔΔCt method to calculate fold change, with *7SK* as the control gene.

### Morpholino, RNA injection and embryological methods

The *ints13* morpholino (MO) purchased from Gene Tools were resuspended in sterile water to a concentration of 1 mM according to the manufacturer’s instructions. RNA was then extracted and converted to cDNA. To target non-neural ectoderm, 1:4 of diluted *ints13*-MO was injected at the two-to four-cell stage. Morpholino sequences are given in Supplemental Table 2. Protocols for *Xenopus* fertilization, MO-injections, antibody staining, whole-mount in situ hybridization and immunofluorescence staining are at our protocol website (https://sites.google.com/a/reversade.com/www/protocols/). All experiments with *Xenopus* embryos were approved by the Singapore National Advisory on Laboratory Animal Research. A stereomicroscope equipped with an ICD digital camera (Leica M205 FA) was used to capture images of embryos in successive focal planes. Images were then combined into one picture with Photoshop CS6 (Adobe).

### Western Blot

For Figures 2 and 6: Cells or embryos were lysed in RIPA buffer supplemented with a cocktail of protease inhibitor (Roche). Total protein was resolved on SDS polyacrylamide gels (Bio-Rad) with DTT, followed by transfer on polyvinylidene difluoride (PVDF) membranes. The following antibodies were used for HRP-mediated chemiluminescent: polyclonal Rabbit antibody to INTS13 (C & M), monoclonal Mouse antibody to actin (Chemicon; Mab1501R), β-tubulin (E7, Developmental Studies Hybridoma Bank, University of Iowa, Iowa City, IA), dynein heavy chain (P1H4, gift from T. Hays, University of Minnesota, Minneapolis, MN) and anti-HA antibody (Sigma, H 6533 (clone HA-7).

For Figures 4 and 5: Protein was extracted directly from RPE cells treated with siRNA by adding 2X SDS loading buffer (120mM Tris pH6.8, 4% SDS, 200mM DTT, 20% Glycerol, and 0.02% Bromophenol blue) to cells while on the plate. Samples were then incubated at room temperature while on the plate with periodic swirling prior to a 10-minute boiling at 95°C and a short sonication. Denatured protein samples were then resolved on a 10% polyacrylamide gel and transferred to a PVDF membrane (Thermo Scientific, 88518). Blots were probed as previously described ^37^. Samples from nuclear extract immunoprecipitation are described in the anti-FLAG affinity purification.

### Electron microscopy (EM)

Embryos at the desired stages were fixed in 3% PBS-glutaraldehyde, pH 7.4, for 4 hours with agitation. After 3 washes in fresh phosphate buffer (NaH2PO4,2H2O / Na2HPO4,12H2O) for 5 minutes each with vigorous agitation, embryos were rinsed 3 times / 5 minutes in distilled water, dehydrated in ethanol with vigorous agitation, dried in Leica EM CPD030 and sputter coated in gold coating, 7nm thick layer using Leica SCD050. Embryos were viewed using a scanning electron microscopy (FESEM, JEOL, JSM-6701F). For Transmission electron microscopy (TEM) dehydrated embryos were embedded in Epon 812, prepared sections (50-60 nm) and contrasted with 4% uranyl acetate in 50% ethanol and 2.6% lead nitrate in 1 M NaOH. Embryos were scanned using a Hitachi 7000 electron microscope (Hitachi, Tokyo, Japan). EM scanning was performed at electron microscopy facility (Institute of Molecular and Cell Biology (IMCB), A*STAR, Singapore).

### Plasmid construction and Yeast two-hybrid analysis

Yeast two-hybrid analysis was carried out in PJ69-4α and PJ49-4a strains as described^33^. In brief, human INTS1 through INTS14, and INTS13 variations, were cloned into pGAD, pOBD, pTEF, or pCEV-G1Ura3/TEFdual vectors using conventional cloning. pOBD plasmids were then transformed into PJ69-4α yeast and were selected on tryptophan-dropout medium (Clontech, Mountain View CA, #630413); pGAD plasmids were transformed into PJ49-4a yeast and were selected on leucine-dropout medium (Clontech, #630414). pTEF or pTEFdual plasmids were included to express proteins in trans and were transformed with pOBD plasmids and selected on media lacking tryptophan and uracil (Sunrise Science Products, San Diego CA, 1316-030). Yeast strains were then mated and subsequently selected on medium lacking tryptophan, leucine, and uracil (Sunrise Science Products, 1328-030). Interactions were tested through serial dilution (one to five) of diploid yeast followed by plating on medium lacking tryptophan, leucine, and uracil or on medium lacking tryptophan, leucine, uracil, and histidine (Sunrise Science Products, 1334-030) that also was supplemented with 1 mM 3-amino-1,2,4-triazole (MP Biomedicals, Irvine CA, 4061-722).

### Expression and purification of INTS4-INTS9-INTS11-INTS13 complex

Full-length human INTS4, INTS9, INTS11, and INTS13 were co-expressed in insect cells using Multibac technology (Geneva Biotech)^71^. INTS4 carried an N-terminal hexa-histidine tag while the other three proteins were untagged. High Five cells were grown in ESF 921 medium (Expression Systems) by shaking at 120 rpm at 27 °C until the density reached 1.5-2×10^6^ cells mL^−1^. Cells were infected by P1 viruses and harvested after 48 h. The cells were lysed by sonication in a buffer containing 20 mM Tris (pH 8.0), 200 mM NaCl, 5% (v/v) glycerol and 0.1 mM PMSF. The supernatant of cell lysates was incubated with nickel beads (Qiagen) and the eluate was then subjected to gel filtration chromatography (Superdex 200, GE Healthcare), in a running buffer of 20 mM Tris (pH 8.0), 250 mM NaCl, and 5 mM DTT.

### Nuclear extract

Cells were collected and washed in cold PBS. 293T cells were detached from dishes while S2 cells were grown in suspension and pelleted by centrifugation. Cells were then resuspended in five times the cell pellet volume of Buffer A (10mM Tris pH8, 1.5 mM MgCl_2_, 10 mM KCl, 0.5mM DTT, and 0.2mM PMSF). Resuspended cells were allowed to swell during a 15-minute rotation at 4°C. After pelleting down at 1,000g for 10 minutes, two volumes of the original cell pellet of Buffer A was added and cells were homogenized with a dounce pestle B for 20 strokes on ice. Nuclear and cytosolic fractions were then separated by centrifugation at 2,000g for 10 minutes. To attain a nuclear fraction, the pellet was washed once with Buffer A before resuspending in an equal amount of the original cell pellet volume of Buffer C (20 mM Tris pH8, 420mM NaCl, 1.5 mM MgCl_2_, 25% glycerol, 0.2 mM EDTA, 0.5 mM PMSF, and 0.5 mM DTT). The sample was then homogenized with a dounce pestle B for 20 strokes on ice and rotated for 30 minutes at 4°C before centrifuging at 15,000g for 30 minutes at 4°C. Finally, supernatants were collected and subjected to dialysis in Buffer D (20 mM HEPES, 100 mM KCl, 0.2 mM EDTA, 0.5 mM DTT, and 20% glycerol) overnight at 4°C. Prior to any downstream applications, nuclear extracts were centrifuged again at 15,000g for 3 minutes at 4°C to remove any precipitate.

### Anti-FLAG affinity purification

To purify FLAG-tagged Integrator complexes for mass spectrometry, generally between 8 and 10 mg of nuclear extract (approximately 1.9 mL of extract depending on the concentration) was mixed with 100 μL anti-Flag M2 affinity agarose slurry (Sigma-Aldrich, #A2220) washed with 0.1 M glycine then equilibrated in binding buffer (20 mM HEPES pH7.4, 150 mM KCl, 10% Glycerol, 0.1% NP-40). This mixture was rotated for four hours at 4°C. Following the four-hour incubation/rotation, five sequential washes were carried out in binding buffer with a 10-minute rotation at 4°C followed by a 1,000g centrifugation at 4°C. After a final wash with 20 mM HEPES buffer, the supernatant was removed using a pipette and the beads were kept cold and submitted to the mass spectrometry core where the protein complexes were eluted by digestion (described below). For immunoprecipitation samples intended for Western blot, a similar protocol was used. 25 µL of bead slurry and 200 µL of extract sample were rotated for two hours at 4°C. After the fifth wash with binding buffer, protein complexes were eluted from the anti-FLAG resin by adding 50 μL of 2X SDS loading buffer and boiled at 95°C for five minutes. For Western blots, input samples were generated by adding equal volume of 2X SDS loading buffer to nuclear extract and 1/10 of the immunoprecipitation was loaded as estimated by protein mass.

### Mass spectrometry sample digestion

The samples were prepared in a similar manner as described previously ^72^. Briefly, the agarose bead-bound proteins were washed several times with 50 mM Triethylammonium bicarbonate (TEAB) pH 7.1, before being solubilized with 40 μL of 5% SDS, 50 mM TEAB, pH 7.55 followed by a room temperature incubation for 30 minutes. The supernatant containing the proteins of interest was then transferred to a new tube, reduced by making the solution 10 mM Tris(2-carboxyethyl)phosphine (TCEP) (Thermo, #77720), and further incubated at 65°C for 10 minutes. The sample was then cooled to room temperature and 1 μL of 1M iodoacetamide acid was added and allowed to react for 20 minutes in the dark. Then, 5 μL of 12% phosphoric acid was added to the 50 μL protein solution followed by 350 μL of binding buffer (90% Methanol, 100 mM TEAB final; pH 7.1). The resulting solution was administered to an S-Trap spin column (Protifi, Farmingdale NY) and passed through the column using a bench top centrifuge (30 second spin at 4,000g). The spin column was then washed three times with 400 μL of binding buffer and centrifuged (1200 rpm, 1 min). Trypsin (Promega, #V5280) was then added to the protein mixture in a ratio of 1:25 in 50 mM TEAB, pH=8, and incubated at 37°C for 4 hours. Peptides were eluted with 80 µL of 50 mM TEAB, followed by 80 μL of 0.2% formic acid, and finally 80 μL of 50% acetonitrile, 0.2% formic acid. The combined peptide solution was then dried in a speed vacuum (room temperature, 1.5 hours) and resuspended in 2% acetonitrile, 0.1% formic acid, 97.9% water and aliquoted into an autosampler vial.

### NanoLC MS/MS Analysis

Peptide mixtures were analyzed by nanoflow liquid chromatography-tandem mass spectrometry (nanoLC-MS/MS) using a nano-LC chromatography system (UltiMate 3000 RSLCnano, Dionex, Thermo Fisher Scientific, San Jose, CA). The nano-LC-MS/MS system was coupled on-line to a Thermo Orbitrap Fusion mass spectrometer (Thermo Fisher Scientific, San Jose, CA) through a nanospray ion source (Thermo Scientific). A trap and elute method was used to desalt and concentrate the sample, while preserving the analytical column. The trap column (Thermo Scientific) was a C18 PepMap100 (300 µm X 5 mm, 5 µm particle size) while the analytical column was an Acclaim PepMap 100 (75 μm X 25 cm) (Thermo Scientific). After equilibrating the column in 98% solvent A (0.1% formic acid in water) and 2% solvent B (0.1% formic acid in acetonitrile (ACN)), the samples (2 µL in solvent A) were injected onto the trap column and subsequently eluted (400 nL/min) by gradient elution onto the C18 column as follows: isocratic at 2% B, 0-5 min; 2% to 32% B, 5-39 min; 32% to 70% B, 39-49 min; 70% to 90% B, 49-50 min; isocratic at 90% B, 50-54 min; 90% to 2%, 54-55 min; and isocratic at 2% B, until the 65 minute mark.

All LC-MS/MS data were acquired using XCalibur, version 2.1.0 (Thermo Fisher Scientific) in positive ion mode using a top speed data-dependent acquisition (DDA) method with a 3 second cycle time. The survey scans (m/z 350-1500) were acquired in the Orbitrap at 120,000 resolution (at m/z = 400) in profile mode, with a maximum injection time of 100 m s and an AGC target of 400,000 ions. The S-lens RF level was set to 60. Isolation was performed in the quadrupole with a 1.6 Da isolation window, and CID MS/MS acquisition was performed in profile mode using rapid scan rate with detection in the ion-trap using the following settings: parent threshold = 5,000; collision energy = 32%; maximum injection time 56 msec; AGC target 500,000 ions. Monoisotopic precursor selection (MIPS) and charge state filtering were on, with charge states 2-6 included. Dynamic exclusion was used to remove selected precursor ions, with a +/-10 ppm mass tolerance, for 15 seconds after acquisition of one MS/MS spectrum.

### Database Searching

Tandem mass spectra were extracted and charge state deconvoluted using Proteome Discoverer (Thermo Fisher, version 2.2.0388). Deisotoping was not performed. All MS/MS spectra were searched against the appropriate database, either Uniprot *Drosophila* database (version 04-04-2018) or Uniprot Human database (reviewed 06272018), using Sequest. Searches were performed with a parent ion tolerance of 5 ppm and a fragment ion tolerance of 0.60 Da. Trypsin was specified as the enzyme, allowing for two missed cleavages. Fixed modification of carbamidomethyl (C) and variable modifications of oxidation (M) and deamidation were specified in Sequest. Heat maps in Figure 4 were made using Morpheus from the Broad Institute, https://software.broadinstitute.org/morpheus.

### RNA interference

siRNA used were Universal Negative Control #2 (Sigma Aldrich, #SIC002), INTS13-1 (Sigma Aldrich, SASI_Hs01_00134940/ASUN), and INTS13-2 (Sigma Aldrich, SASI_Hs01_00134941/ASUN). RPE cells were seeded at 50,000 cells/well in a 24-well plate on Day 1. On Day 2, 3 µL of 20 µM siRNA was added to 50 µL of Opti-MEM I per well and incubated at room temperature for 5 minutes. 2 µL of Lipofectamine RNAiMax (Invitrogen, 13778-150) was also added to 50 µL of Opti-MEM per well and incubated at room temperature for 5 minutes. The siRNA and RNAi Max solutions were combined and incubated at room temperature for 20 minutes, then added to the cells. Cells were expanded to a 6-well plate on Day 3. On Day 4, cells were washed gently with DPBS and 900 µL of DMEM/F-12 media was added to each well. The siRNA treatment from Day 2 was repeated. Cells were further incubated and then harvested on Day 6. For RNA extraction, cells were harvested with 500 µL of Trizol. For protein extraction, cells were harvested with SDS loading buffer detailed previously.

A modified version of the above protocol was followed for cells intended for immunofluorescence staining of primary cilia: RPE cells were seeded onto three cover slips in each well of the 6-well plate on Day 3. On Day 4, cells were washed twice with DPBS and grown in serum free (0% FBS) DMEM/F-12 + 1% (v/v) penicillin-streptomycin for 48 hours before fixation.

### Library preparation and RNA sequencing

RNAi-treated RPE cells were harvested in 500 µL of Trizol, and total RNA was purified by phenol-chloroform extraction. 1 µg total RNA per sample was used for sequencing library preparation. The NEBNext Poly(A) mRNA magnetic isolation kit (New England Biolabs, Ipswich MA, E7490) was used to purify mRNA from the total RNA. The Click-Seq protocol was used following this step^73^. RT-PCR to make cDNA was done with 1:35 azido-NTP:dNTP for random strand termination by SuperScript III reverse transcriptase. Following Click-Seq preparations, libraries were PCR amplified for 17 cycles and products from 400-600 bp were collected for sequencing. Samples were sequenced on a NextSeq500 (Illumina) using a single read 75bp run. Sequencing adapters removed and reads with a Qscore<30 were discarded using fastp (ver 0.14.1). Reads were aligned using Hisat2 (ver 2.1.0) to the hg38 genome using default parameters. Differential expression changes were determined using DESeq2 (ver 1.23.10) and assigned using the comprehensive gene annotation in GENCODE32 and tests for significance were using the nbinomialWald test. Alignment information and lowest correlation between samples in each data set is given in table below.

| Sample | Total Reads | Mappable Fragments (Percent of total) | Agreement between replicates (Spearman's rho) | Normalization Factor |
| --- | --- | --- | --- | --- |
| Control | 131,005,450 | 94.33% | >0.99 | 1.142296033 |
| INTS13-1 | 149,803,846 | 94.27% | >0.99 | 0.999334293 |
| INTS13-2 | 168,253,066 | 94.13% | >0.99 | 0.889755677 |

## ACKNOWLEDGMENTS

We thank all families for partaking in this study. The authors would also like to thank members of their team for technical assistance and fruitful discussions. This work was supported by a grant from the French Ministry of Health to J.A., Strategic Positioning Fund on Genetic Orphan Diseases from the Biomedical Research Council, A*STAR, Singapore to B.R., and by the UCLA California Center for Rare Diseases to H.L., S.F.N. B.R. is a fellow of the Branco Weiss Foundation, an A*STAR Investigator and Young EMBO Investigator. This work was supported by National Institutes of Health grant R01-GM134539 (E.J.W.), and the Welch Foundation grant H-1889 to The University of Texas Medical Branch at Galveston (E.J.W.). The UTMB Mass Spectrometry Facility is supported in part by The Cancer Prevention Research Institute of Texas (CPRIT) grant number RP190682 (W.K.R). M.K.S. was supported by CPRIT grant number P71709-B01.

## AUTHOR CONTRIBUTIONS

L.G.M. and M.S. contributed equally to this work. B.R. initiated this project with H.H. who clinically ascertained Family 1 with the support of B.A.R., O.A. and M.E.K. Family 2 was ascertained by J.A. and L.C. L.G.M. conducted RPE analysis, biochemical purifications, and yeast two-hybrid analysis. M.S. performed all experiments on patients’ primary cells and in *Xenopus* with the help of N.E.B. N.D.E. prepared samples for and analyzed RNA-seq. K-L.H. performed Western blot analysis. N.P. built the pTEFdual vector. H.L., S.F.N., and B.M. conducted linkage/sequence analysis. Y.X. and L.T. conducted recombinant protein production and gel filtration. M.K.S. and L.J.K. imaged and analyzed PC immunofluorescence. W.K.R. prepared proteomic samples and analyzed LC/MS data. L.G.M., M.S., B.R., and E.J.W. wrote the manuscript. All authors critically reviewed the manuscript.

## WEB RESOURCES

The URLs for data presented herein are as follows:

BLAST, http://blast.ncbi.nlm.nih.gov/Blast.cgi

dbSNP146, http://www.ncbi.nlm.nih.gov/snp

1000 Genomes, http://www.1000genomes.org/

Online Mendelian Inheritance in Man (OMIM), http://www/omim.org/

ExAC, http://exac.broadinstitute.org/

ANNOVAR, http://www.openbioinformatics.org/annovar/

PolyPhen-2, http://genetics.bwh.harvard.edu/pph2/

Human Splicing Finder, http://www.umd.be/HSF3/HSF.html

ClinVar, ftp://ftp.ncbi.nlm.nih.gov/pub/clinvar/vcf_GRCh37/

Exome Sequencing Project, http://evs.gs.washington.edu/EVS/

## SUPPLEMENTAL FIGURE LEGENDS

**Supplemental Fig 1. Presentation of OFD2 Syndrome patients from Family 1 and Family 2. a-d, m** Photographs of affected individual II.4 of Family 1 with common phenotype and orofacial dysmorphisms at different ages including: **a-c** Bilateral cleft lip and palate (corrected after multiple surgical operations), hypertelorism, broad nasal bridge, flat philtrum, dental abnormalities, low set ear and rough and sparse hair. **d** Crowded optic disc. **m** Clinodactyly of toes. **e-l** Photographs of affected individual II.5 of Family 1. **e-g** Bilateral cleft lip and palate, hypertelorism, broad nasal bridge, flat philtrum, dental abnormalities. **h** Crowded optic disc. **i** Low set ear and rough and sparse hair. **j, h** Broad and mild brachydactyly and single palmar creases. **l** Clinodactyly of toes. **n** Pedigree of Family I. The genotypes for the corresponding mutation are indicated below each individual. The c.2004delA mutation segregated with the disease in this family. (+/-) denotes the heterozygous alleles, and (-/-) denotes the homozygous mutant alleles. **o-v** Photographs of affected individuals II.2 and II.4 of Family 2 at 27 and 20 years old, respectively. **w** Pedigree of Family 2. The genotypes for the corresponding mutation are shown below each individual which confirm the segregation of the c.1955C>T mutation in this family. (+/+) denotes the homozygous wildtype alleles.

**Supplemental Fig 2. The C-terminus of INTS13 interacts with the Integrator Cleavage Module. a** Expanded modified yeast two hybrid assay from Fig 2a. Included here are tests where INTS11 or INTS9 were expressed in *trans* along with INTS13-binding domain and crossed with each Integrator subunit. No positive interactions were seen. **b** Left: Modified yeast two hybrid with INTS9 fused to the binding domain and INTS11 expressed in *trans* mated with yeast expressing each Integrator subunit fused to the activating domain to reconfirm that INTS4 interacts with the INTS9/11 heterodimer. Right: The same experiment with the addition of INTS4 expressed in *trans* to show that INTS13 interacts with the INTS4/9/11 heterotrimer. **c** Smaller portions (approximately 20 amino acids) of the INTS13 protein were removed from the necessary region for interaction shown in Fig 2d to determine that residues 577-706 were sufficient to interact with INTS4/9/11.

**Supplemental Fig 3. INTS13 interacts with INTS10/14. a** Partial protein alignment of the C-terminus of human and *Drosophila* INTS13 to show the orthologous mutations made to recapitulate the patient mutations in *Drosophila* cells. The location and altered residues are marked in brown for Family 1 and in green for Family 2. **b** Modified yeast two hybrid testing the interaction between INTS10/13/14. Left: INTS14 fused to the binding domain was crossed with each Integrator subunit fused to the activating domain, and no positive interaction was seen on selective media. Right: INTS13 fused to the activating domain was crossed with yeast strains expressing INTS14-binding domain with each Integrator subunit in *trans*. A positive interaction was seen when INTS10 was expressed in *trans*.

**Supplemental Fig 4. Significant overlap in gene expression changes between two *INTS13*-targeting siRNAs. a** Venn diagrams show the number of significantly upregulated or **b** downregulated genes for siRNA 13-1 and 13-2, and the genes in common between the two siRNAs as determined by RNA-seq.

**Supplemental Fig 5. Expression of *ints13* during *Xenopus* development. a** Detailed expression pattern of *ints13* in *Xenopus* embryos analyzed by WISH and whole mount immunostaining. *ints13* has both maternal and zygotic contribution. At 4 cell stage, *ints13* transcripts are localized within the animal half. At stage 10, its expression covers the whole embryo except the blastopore. At stage 17, *ints13* is observed in neural tube and neural folds. **b** Lateral view, anterior is right stage 22 *ints13* starts to be enriched in brain, branchial arches (ba), pronephros (pn), eye vesicles (ev), ear (e), skin cells (sk) and somites (so). Lateral view, anterior is right of the embryo at stage 28, *ints13* expression is also detected in ear (e), skin cells (sk) and pronephros (pn). Correlation of transcripts localization of ints13 protein at stage 28 by wholemount embryo antibody staining using a custom polyclonal INTS13 (α-M) antibody, Lateral view, anterior is left.

**Table 1.**
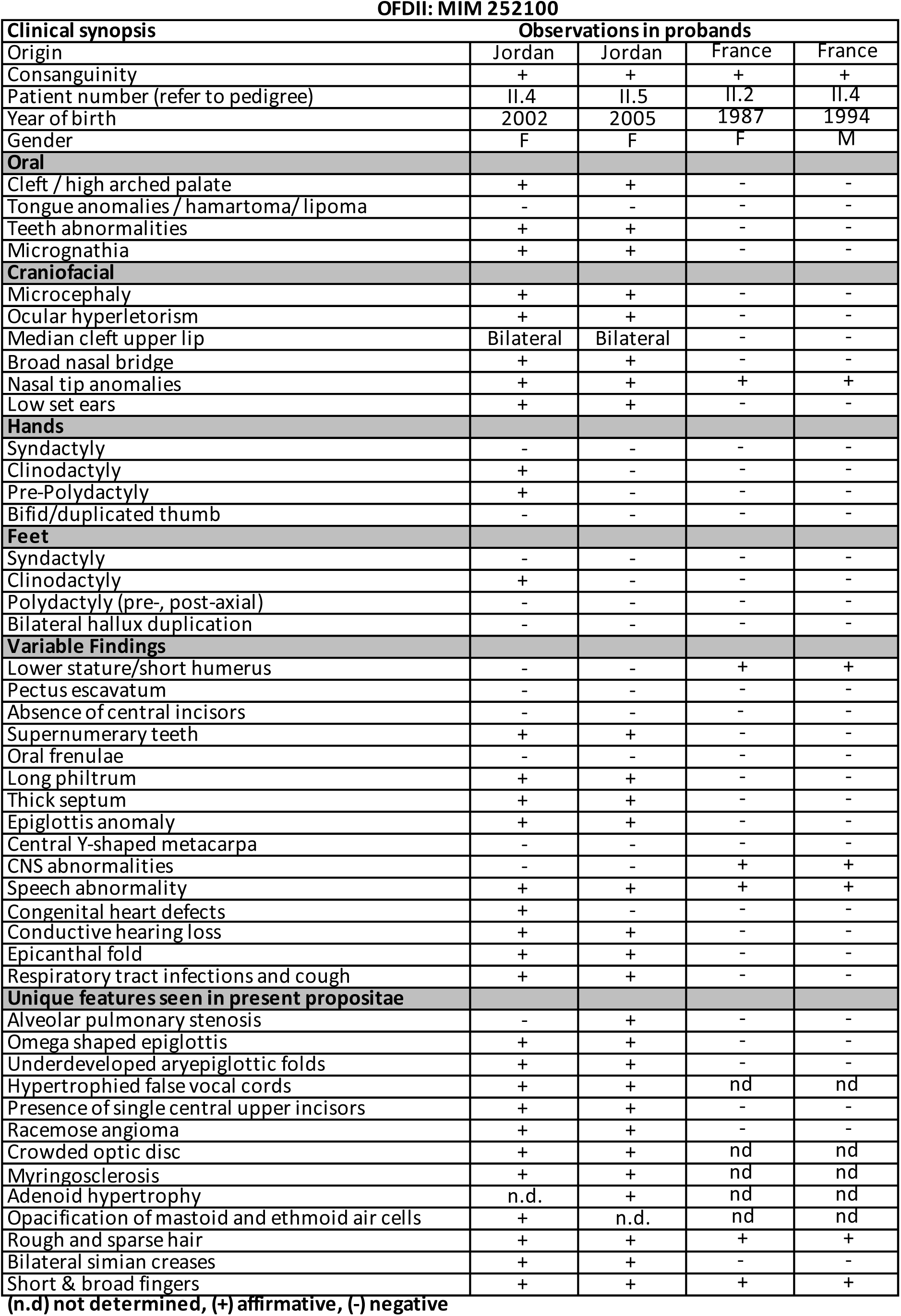
Clinical phenotypes of the affected individuals. (n.d) not determined, (+) affirmative, (-) negative, (M) male and (F) female.

**OFDII: MIM 252100**
| **Clinical synopsis** |  | **Observations in probands** |  |  |  |
| --- | --- | --- | --- | --- | --- |
| Origin |  | Jordan | Jordan | France | France |
| Consanguinity |  | + | + | + | + |
| Patient number (refer to pedigree) |  | II.4 | II.5 | II.2 | II.4 |
| Year of birth |  | 2002 | 2005 | 1987 | 1994 |
| Gender |  | F | F | F | M |
| **Oral** |  |  |  |  |  |
| Cleft / high arched palate |  | + | + | - | - |
| Tongue anomalies / hamartoma / lipoma |  | - | - | - | - |
| Teeth abnormalities |  | + | + | - | - |
| Micrognathia |  | + | + | - | - |
| **Craniofacial** |  |  |  |  |  |
| Microcephaly |  | + | + | - | - |
| Ocular hyperletorism |  | + | + | - | - |
| Median cleft upper lip |  | Bilateral | Bilateral | - | - |
| Broad nasal bridge |  | + | + | - | - |
| Nasal tip anomalies |  | + | + | + | + |
| Low set ears |  | + | + | - | - |
| **Hands** |  |  |  |  |  |
| Syndactyly |  | - | - | - | - |
| Clinodactyly |  | + | - | - | - |
| Pre-Polydactyly |  | + | - | - | - |
| Bifid/duplicated thumb |  | - | - | - | - |
| **Feet** |  |  |  |  |  |
| Syndactyly |  | - | - | - | - |
| Clinodactyly |  | + | - | - | - |
| Polydactyly (pre-, post-axial) |  | - | - | - | - |
| Bilateral hallux duplication |  | - | - | - | - |
| **Variable Findings** |  |  |  |  |  |
| Lower stature/short humerus |  | - | - | + | + |
| Pectus excavatum |  | - | - | - | - |
| Absence of central incisors |  | - | - | - | - |
| Supernumerary teeth |  | + | + | - | - |
| Oral frenulae |  | - | - | - | - |
| Long philtrum |  | + | + | - | - |
| Thick septum |  | + | + | - | - |
| Epiglottis anomaly |  | + | + | - | - |
| Central Y-shaped metacarpa |  | - | - | - | - |
| CNS abnormalities |  | - | - | + | + |
| Speech abnormality |  | + | + | + | + |
| Congenital heart defects |  | + | - | - | - |
| Conductive hearing loss |  | + | + | - | - |
| Epicanthal fold |  | + | + | - | - |
| Respiratory tract infections and cough |  | + | + | - | - |
| **Unique features seen in present propositae** |  |  |  |  |  |
| Alveolar pulmonary stenosis |  | - | + | - | - |
| Omega shaped epiglottis |  | + | + | - | - |
| Underdeveloped aryepiglottic folds |  | + | + | - | - |
| Hypertrophied false vocal cords |  | + | + | nd | nd |
| Presence of single central upper incisors |  | + | + | - | - |
| Racemose angioma |  | + | + | - | - |
| Crowded optic disc |  | + | + | nd | nd |
| Myringosclerosis |  | + | + | nd | nd |
| Adenoid hypertrophy |  | n.d. | + | nd | nd |
| Opacification of mastoid and ethmoid air cells |  | + | n.d. | nd | nd |
| Rough and sparse hair |  | + | + | + | + |
| Bilateral simian creases |  | + | + | - | - |
| Short & broad fingers |  | + | + | + | + |

**Supplemental Table 1. RNA-seq differential expression analysis results**.

**Supplemental Table 2. Primer sequences**.

