## Supplemental Figure 1 for "*INTS13* Mutations Causing a Developmental Ciliopathy Disrupt Integrator Complex Assembly"

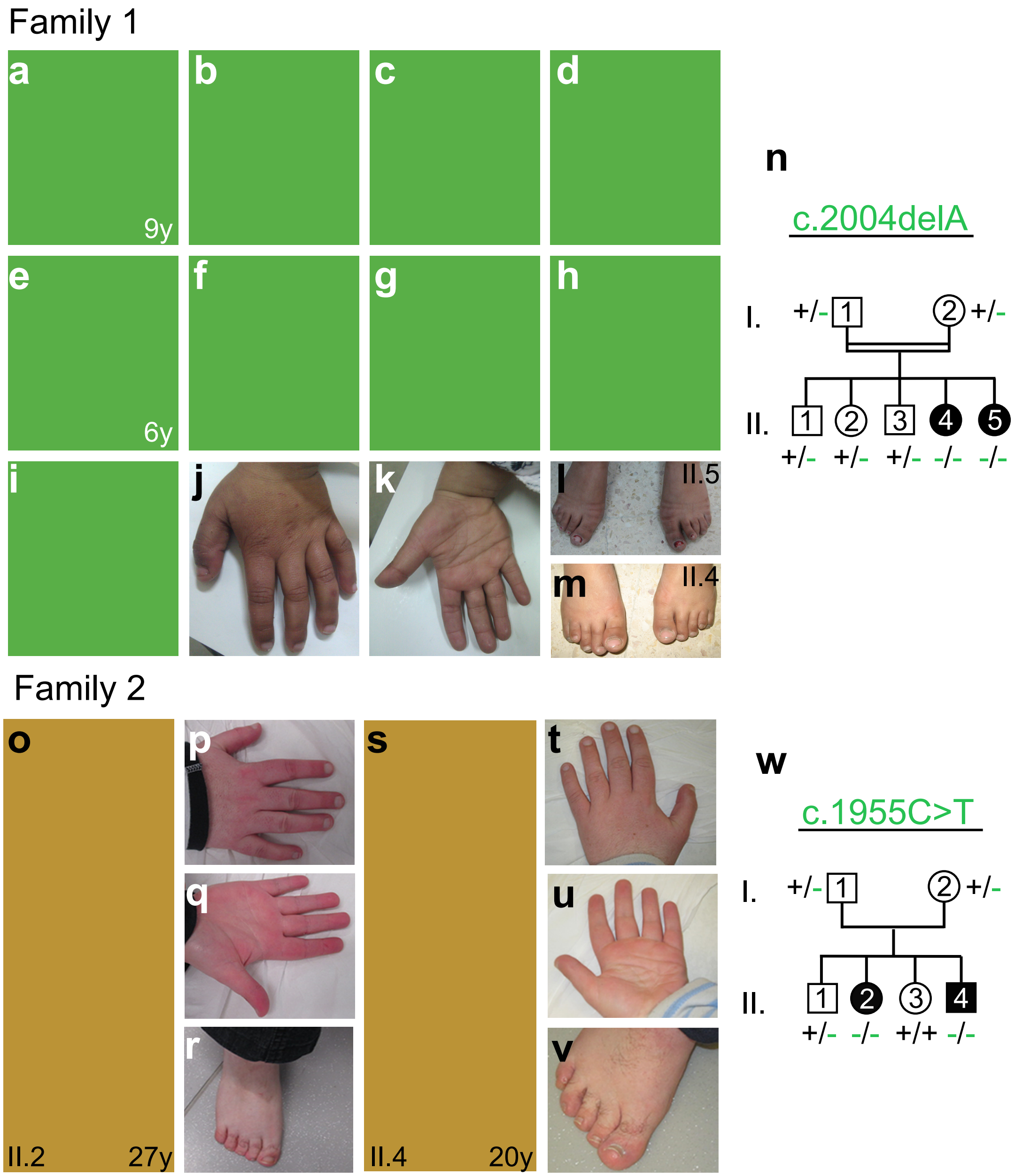


**Supplemental Fig 1.** Presentation of OFD2 Syndrome patients from Family 1 and Family 2. **a-d, m** Photographs of affected individual II.4 of Family 1 with common phenotype and orofacial dysmorphisms at different ages including: **a-c** Bilateral cleft lip and palate (corrected after multiple surgical operations), hypertelorism, broad nasal bridge, flat philtrum, dental abnormalities, low set ear and rough and sparse hair. **d** Crowded optic disc. m Clinodactyly of toes. **e-l** Photographs of affected individual II.5 of Family 1. **e-g** Bilateral cleft lip and palate, hypertelorism, broad nasal bridge, flat philtrum, dental abnormalities. **h** Crowded optic disc. **i** Low set ear and rough and sparse hair. **j, h** Broad and mild brachydactyly and single palmar creases. **l** Clinodactyly of toes. n Pedigree of Family I. The genotypes for the corresponding mutation are indicated below each individual. The c.2004delA mutation segregated with the disease in this family. (+/-) denotes the heterozygous alleles, and (-/-) denotes the homozygous mutant alleles. **o-v** Photographs of affected individuals II.2 and II.4 of Family 2 at 27 and 20 years old, respecitvely. **w** Pedigree of Family 2. The genotypes for the corresponding mutation are shown below each individual which confirm the segregation of the c.1955C>T mutation in this family. (+/+) denotes the homozygous wildtype alleles.
