## Supplemental Figure 2 for "*INTS13* Mutations Causing a Developmental Ciliopathy Disrupt Integrator Complex Assembly"

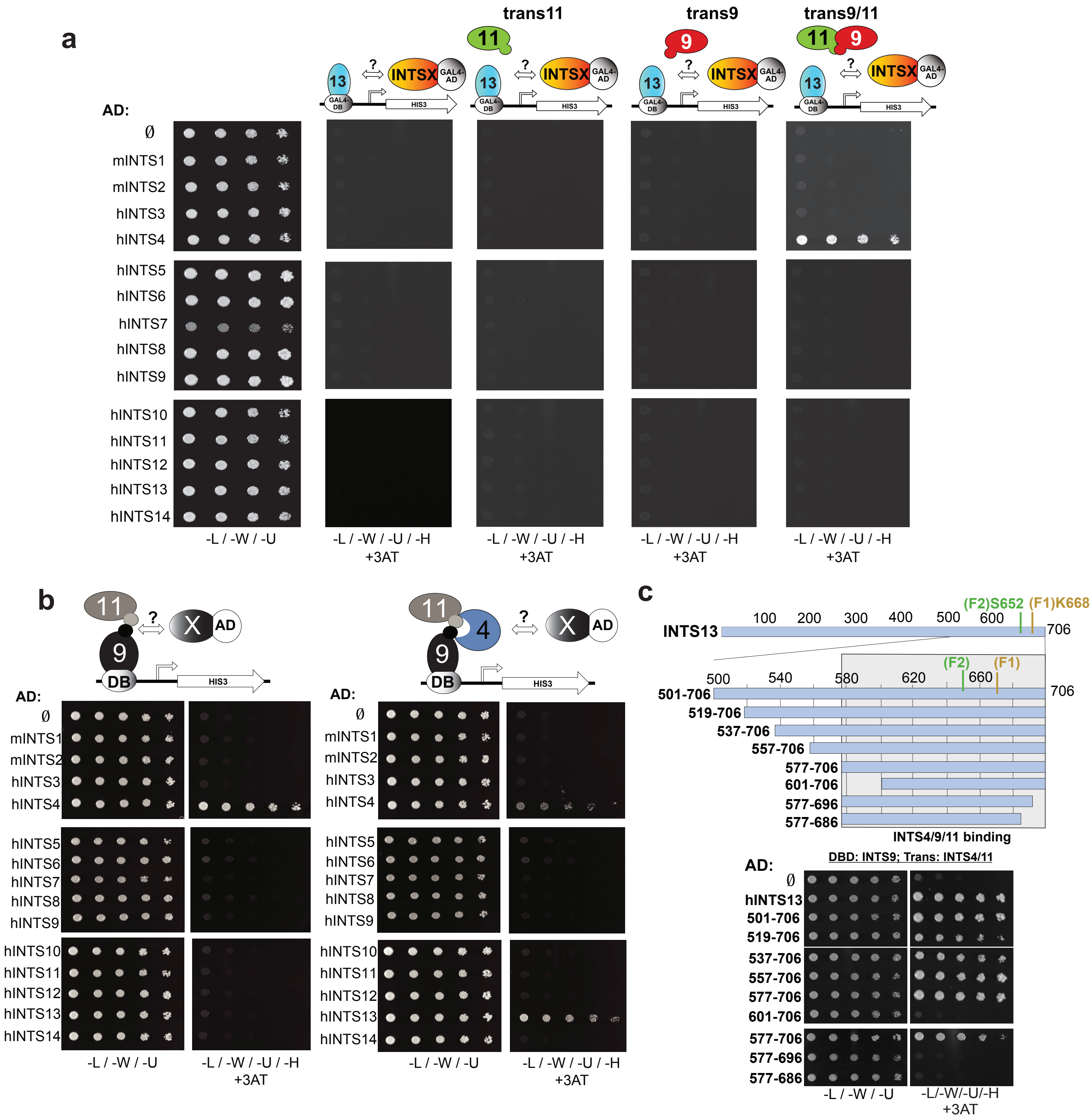


**Supplemental Fig 2. The C-terminus of INTS13 interacts with the Integrator Cleavage Module. a** Expanded modified yeast two hybrid assay from Fig 2a. Included here are tests where INTS11 or INTS9 were expressed in trans along with INTS13-binding domain and crossed with each Integrator subunit. No positive interactions were seen. **b** Left: Modified yeast two hybrid with INTS9 fused to the binding domain and INTS11 expressed in trans mated with yeast expressing each Integrator subunit fused to the activating domain to reconfirm that INTS4 interacts with the INTS9/11 heterodimer. Right: The same experiment with the addition of INTS4 expressed in trans to show that INTS13 interacts with the INTS4/9/11 heterotrimer. **c** Smaller portions (approximately 20 amino acids) of the INTS13 protein were removed from the necessary region for interaction shown in Fig 2d to determine that residues 577-706 were sufficient to interact with INTS4/9/11.
