## Supplemental Figure 3 for "*INTS13* Mutations Causing a Developmental Ciliopathy Disrupt Integrator Complex Assembly"

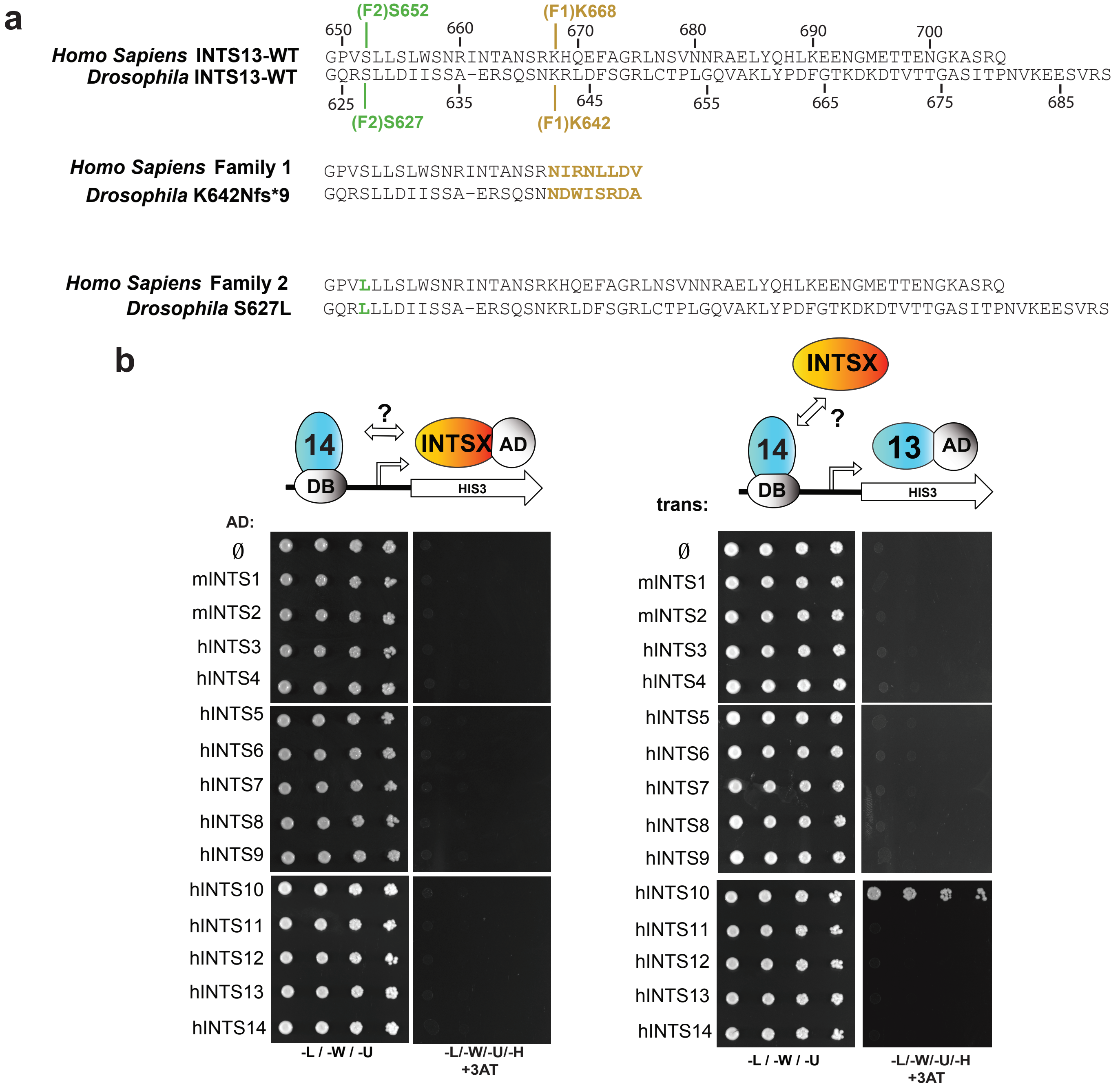


**Supplemental Fig 3**. INTS13 interacts with INTS10/14. **a** Partial protein alignment of the C-terminus of human and Drosophila INTS13 to show the orthologous mutations made to recapitulate the patient mutations in Drosophila cells. The location and altered residues are marked in brown for Family 1 and in green for Family 2. **b** Modified yeast two hybrid testing the interaction between INTS10/13/14. Left: INTS14 fused to the binding domain was crossed with each Integrator subunit fused to the activating domain, and no positive interaction was seen on selective media. Right: INTS13 fused to the activating domain was crossed with yeast strains expressing INTS14-binding domain with each Integrator subunit in trans. A positive interaction was seen when INTS10 was expressed in trans.
