## Supplemental Figure 4 for "*INTS13* Mutations Causing a Developmental Ciliopathy Disrupt Integrator Complex Assembly"

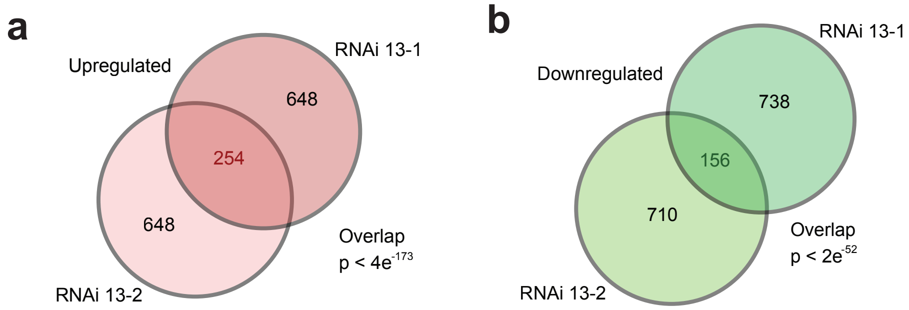


**Supplemental Fig 4. Significant overlap in gene expression changes between two INTS13-targeting siRNAs. a** Venn diagrams show the number of significantly upregulated or **b** downregulated genes for siRNA 13-1 and 13-2, and the genes in common between the two siRNAs as determined by RNA-seq.
