## Supplemental Figure 5 for "*INTS13* Mutations Causing a Developmental Ciliopathy Disrupt Integrator Complex Assembly"

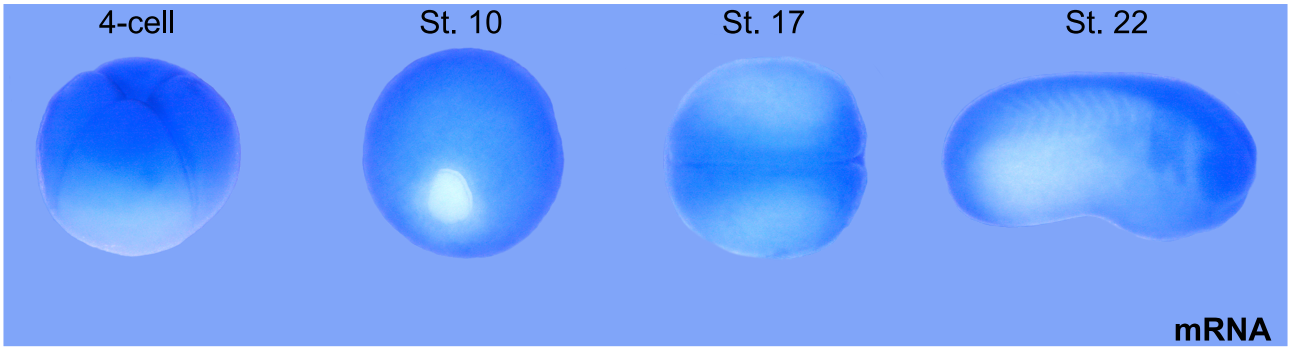


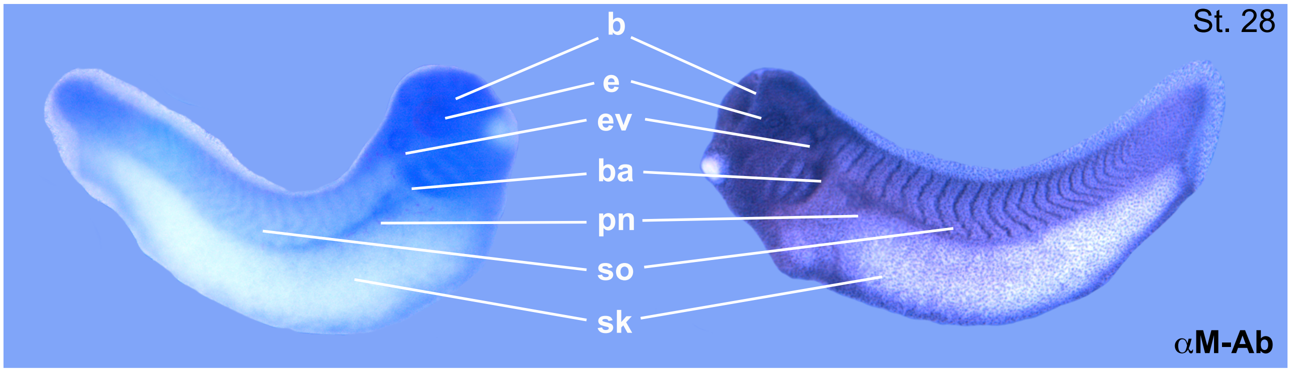


**Supplemental Fig 5. Expression of ints13 during Xenopus development. a** Detailed expression pattern of ints13 in Xenopus embryos analyzed by WISH and whole mount immunostaining. ints13 has both maternal and zygotic contribution. At 4 cell stage, ints13 transcripts are localized within the animal half. At stage 10, its expression covers the whole embryo except the blastopore. At stage 17, ints13 is observed in neural tube and neural folds. b Lateral view, anterior is right stage 22 ints13 starts to be enriched in brain, branchial arches (ba), pronephros (pn), eye vesicles (ev), ear (e), skin cells (sk) and somites (so). Lateral view, anterior is right of the embryo at stage 28, ints13 expression is also detected in ear (e), skin cells (sk) and pronephros (pn). Correlation of transcripts localization of ints13 protein at stage 28 by wholemount embryo antibody staining using a custom polyclonal INTS13 (α-M) antibody, Lateral view, anterior is left.
